# A Genome-wide Visual Screen Identifies Lysophosphatidylcholine as Counter Spatial Regulator of DAG and Sterols in Yeast

**DOI:** 10.64898/2026.01.27.702077

**Authors:** A Henderson, A Lalani, S Ganesan, H Mesa Galloso, N Zung, P Portela, ML Sosa Ponce, K Baetz, M Schuldiner, V Zaremberg

## Abstract

Membrane lipids are heterogeneously distributed across the bilayers of cellular membranes. Cytosolic-facing pools of diacylglycerol (DAG) in the yeast *Saccharomyces cerevisiae* are enriched at both ends of the endomembrane system from the vacuolar membrane to the polarized plasma membrane (PM) of buds. However, how this distribution is maintained remains unknown. Using a genome-wide DAG biosensor screen in yeast, we identify regulators of DAG spatial distribution, enriched in proteins involved in vesicle or lipid transport and in phospholipid or sterol metabolism. A subset of mutants exhibited DAG mislocalization predominantly to the PM, with the most severe phenotype linked to a mutant of a predicted lipase we have named Drl1 (<u>D</u>AG redistribution lipase 1). Reversion of this phenotype required both enzymatic activity and the presence of an intrinsically disordered carboxy-terminal domain. Lipidomic analysis revealed that loss of Drl1 increased cellular lysophosphatidylcholine (LysoPC) levels. Remarkably, we find that supplementing cells with a non-metabolizable LysoPC analogue replicated the mutant DAG phenotype, implicating LysoPC as a novel spatial regulator of DAG. High-resolution imaging suggests that LysoPC reduces the PM sterol pool resulting in DAG expansion into new PM territories. More globally, our work expands the known interplay between various lipids and their co-regulation to maintain accurate membrane properties.

## INTRODUCTION

Most eukaryotic membrane lipids are synthesized in the endoplasmic reticulum (ER) and are then transported to other cellular membranes in a leaflet specific composition, reaching the highest asymmetric lipid distribution between external and cytosolic leaflets at the plasma membrane (PM) (Fairn et al., 2011; van Meer et al., 2008). Unlike membrane lipids with phospho-head groups that require active flipping to accumulate on specific membrane leaflets, sterols, ceramides and diacylglycerols (DAGs) can translocate across membranes; a biological process known as flip-flop (Bennett & Tieleman, 2012). However, we currently lack a clear understanding of the forces that govern their movement as well as their cellular localization in live cells. It has been determined that the flip-flop rate of these three lipids depends on the lipid environment (Devaux et al., 2008).

DAG was the first lipid second messenger to be discovered in the canonical activation pathway of protein kinase C (PKC). Despite a robust understanding of DAG’s contribution to cell signaling, the mechanisms regulating the spatial distribution of DAG within the cell, and across leaflets, remain poorly understood, necessitating methods to track its different cellular pools. We have previously developed a fluorescent DAG probe based on the C1 domain of PKCδ to monitor cytoplasmic facing pools of DAG in budding yeast using fluorescence live microscopy (Ganesan et al., 2016). Two pools of DAG were identified in both the vacuole (equivalent to mammalian lysosome) and at sites of polarized growth (Ganesan et al., 2019). Directed studies using this probe in yeast cells undergoing growth resumption from stationary phase have pointed to triacylglycerol lipolysis for the emergence of a vacuolar DAG pool and *de novo* synthesis of phosphatidylserine (PS) by Cho1 for its transport to sites of polarized growth at the PM of buds (Ganesan et al., 2020). However, despite these initial observations, a clear understanding of how DAG distribution between leaflets is regulated was lacking.

In this study, we conducted an unbiassed genome-wide, high-throughput imaging screen surveying single knockout and hypomorphic yeast collections expressing the DAG probe to identify factors and pathways regulating DAG distribution. From the ⁓6,000 strains imaged, we identified a discrete group of 51 mutants with abnormal DAG distribution. The impacted strains were enriched in genes affecting lipid metabolism and transport, as well as membrane remodeling and cargo delivery pathways. A mutant producing a truncated version of an uncharacterized putative lipase displayed a strong phenotype of DAG localized exclusively to the cell periphery. We determined that the abnormal PM DAG phenotype depended on both the activity of the enzyme and its carboxy end. We therefore named this protein Drl1 (for **D**AG **r**edistribution **l**ipase 1). Lipidomic analysis revealed that ablation of Drl1 increased cellular levels of Lysophosphatidylcholine (LysoPC). Remarkably, addition of a non-metabolizable LysoPC analogue to wild-type (WT) yeast mimicked the strong DAG phenotype displayed by lack of Drl1, suggesting a causal relationship between LysoPC levels and DAG localization. High-resolution live microscopy of WT yeast using fluorescent probes revealed polarized DAG and sterol microdomains coexisting at the PM of buds. Addition of the non-metabolizable LysoPC rapidly dissipated the sterol domains with a concomitant expansion of DAG into new areas of the PM. Collectively, these data support LysoPC as a counter spatial regulator of DAG and sterols in yeast. More globally, it gives new mechanistic insight into the interplay of membrane lipids in maintaining essential PM biophysical properties by modulating mobile lipid pools in response to compositional imbalance.

## RESULTS

### Visual genetic screen identifies vesicular transport and lipid metabolic pathways as DAG spatial regulators

To identify factors and pathways that regulate DAG movement and spatial distribution in yeast, a genome-wide microscopy screen was devised. A previously characterized DAG sensor based on the tandem C1A-C1B domains of mammalian PKCδ (C1δ-GFP DAG) (Ohashi et al., 2009; Sánchez-Bautista et al., 2009; Stahelin et al., 2004, 2005) was used to monitor cytosolic facing DAG pools. In a WT background, this sensor localizes predominantly to the vacuolar membrane as evident by colocalization with the endolysosomal dye FM4-64 (Vida & Emr, 1995), as well as at the PM of buds (Figure 1A)(Ganesan et al., 2019).

**Figure 1.**
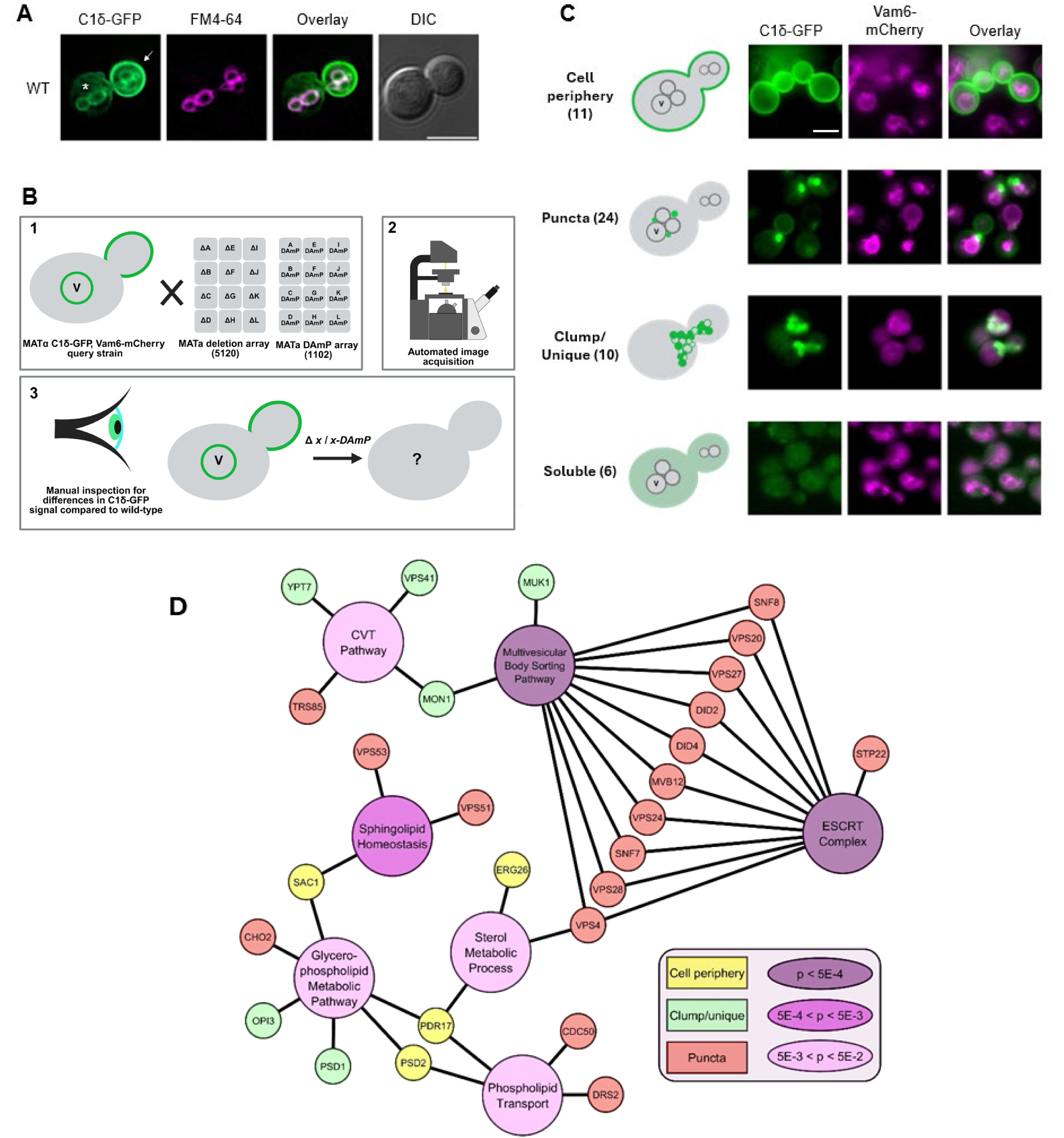
High throughput screen reveals genes involved in DAG localization. **A)** WT cells (*BY4741)* expressing the C1δ-GFP DAG probe were grown to mid-log phase and then incubated with 32 µM FM4-64 for 45 minutes to visualize vacuolar membranes. **(↓)** denotes plasma membrane DAG pool; **(*)** denotes vacuolar DAG pool. Scale bar = 5 µm**B)** Schematic outlining high-throughput microscopy screen. A query strain carrying a plasmid for expression of the C1δ-GFP DAG probe, and an endogenously tagged Vam6-mCherry vacuolar marker was crossed with yeast knockout and DAmP arrays. Following sporulation and selection of haploids, the collection was analyzed using an automated image acquisition system. The images were then manually inspected for strains with aberrant C1δ-GFP signal. **C)** Observed phenotypic categories (cell periphery, puncta, clumps, soluble). The number of mutations linked to each phenotype is shown in parenthesis. Scale bar 5 µm. **D)** Enrichment of genes in our dataset clustering according to Gene Ontology Biological Process and Cellular Component categories for mutants with altered DAG localization using the Cytoscape plugin ClueGo. Phospholipid Transport (GO:0016236), Sterol Metabolic Process (GO:0006886), Glycerophospholipid Metabolic Pathway (GO:0006892), Sphingolipid Homeostasis (GO:0019941), CVT Pathway (GO:0008202), Multivesicular Body Sorting Pathway (GO:0045324), ESCRT Complex (GO:0036452). Enrichment is calculated relative to the frequency of that cluster in the whole genome. Significance of the category is indicated by the node color.

A query strain expressing the C1δ-GFP DAG biosensor from a plasmid, as well as a genomically integrated vacuolar marker (Vam6-RFP) was crossed to two yeast collections using Synthetic Genetic Array (SGA) methodology (Baryshnikova et al., 2010; Cohen & Schuldiner, 2011; Tong et al., 2001)(Figure 1A). For non-essential genes, a knockout collection of 5120 strains was used (Giaever et al., 2002). For essential genes, a hypomorphic collection of 1102 strains using <u>D</u>ecreased <u>A</u>bundance by <u>m</u>RNA <u>p</u>erturbation (DAmP) (Breslow et al., 2008) was employed. This facilitated nearly complete coverage of the yeast genome in a single screen. The resulting haploid yeast library was then subjected to automated image acquisition where three images were acquired for each mutant during mid-logarithmic phase growth. The ∼18,000 images were manually assessed for DAG localization phenotypes which differed from the query WT yeast expressing the C1δ-GFP probe (Figure 1B). From the ⁓6,000 strains imaged, 51 mutants displayed drastic alterations in DAG localization. They were organized in four phenotypic categories: 1) cell periphery, 2) puncta, 3) clumps and 4) soluble (Figure 1C and Supplementary Table 1).

Gene Ontology Biological Process and Cellular Component analyses for the mutants identified in our screen show that genes implicated in vesicular trafficking and lipid transport and homeostasis were highly enriched (Figure 1D). Among the 51 mutants with altered DAG sensor localization, nearly half (24/51, ∼47%) displayed intracellular punctate accumulation of the C1δ-GFP signal (Figure 1C). This “puncta” phenotype mapped largely to genes involved in membrane remodeling and cargo delivery trafficking pathways. Specifically, we identified components of the Golgi-associated retrograde protein (GARP) complex (Vps51, Vps53), the transport protein particle (TRAPP)-III complex (Trs85), and the endosomal sorting complexes required for transport (ESCRT)-III complex (Did4, Did2). Mutants of the lipid flippase Drs2 and its Cdc50 subunit were also represented in this category, pointing to membrane asymmetry as a contributor to DAG localization, in agreement with previously reported results (Ganesan et al., 2019). The second largest category grouped mutants that displayed the DAG sensor at the cell periphery in a non-polarized fashion (11/51, ∼21%) comprising mutants related to lipid metabolism and transport including the C-3 sterol dehydrogenase Erg26, the phosphatidylinositol-4-phosphate (PI4P) phosphatase Sac1, the PS decarboxylase of Golgi and vacuolar membranes Psd2, and its activator the phosphatidylinositol (PI) transfer protein Pdr17. Defining the “clumps/unique” phenotypic category (10/51, ∼20%) were mutants *mon1*Δ, *muk1*Δ and *ypt7*Δ, related to the cytoplasm-to-vacuole targeting (CVT) and multivesicular body (MVB) pathways. Interestingly, cells lacking the other yeast PS decarboxylase, Psd1, which produces phosphatidylethanolamine (PE) in mitochondria, and the PE methyltransferase Opi3 that catalyzes a step in the methylation pathway converting PE to phosphatidylcholine (PC) were among the mutants in this group. The “soluble” category (6/51, ∼12%) displaying diffuse cytosolic localization of the C1δ-GFP sensor, was observed in cells lacking Atp4, Chs5, or Gpx1. Enrichment analysis of all 51 hits identified an interconnected network including the GO cellular categories related to CVT, MVB and ESCRT pathways as well as glycerophospholipid and sterol metabolism and transport (Figure 1D).

### A previously uncharacterized lipase is a strong DAG spatial regulator

We next focused on the “cell periphery” phenotypic category which hosted various uncharacterized genes including *YDR445c*, *YDL085c* and *YDR327w* (Table S1 & Figure S1). *YDR445c* was of particular interest because unlike other hits in this group, the deletion mutant was completely devoid of C1δ-GFP signal in the vacuolar membrane (Figure 2A). Initial inspection of this gene was puzzling as *YDR445c* was annotated as a dubious open reading frame in SGD (Engel et al., 2024). Further inspection revealed that this gene overlapped by 171 base-pairs (bp) with the uncharacterized putative lipase, *YDR444w*. (Figure 2A). Since *YDR444w* was not represented in our tailored SGA collection, we acquired its knockout strain and manually tested DAG distribution. Indeed, *ydr444w*Δ displayed peripheral localization of the DAG sensor as observed in *ydr445c*Δ cells (Figure 2A). Given that the uncharacterized putative lipase, *YDR444w,* exhibited the most consistent and extreme phenotype within a category enriched in genes related to lipid metabolism, we set out to characterize it. Absence of *YDR444w* resulted in a drastic reorganization of cytosolic facing DAG pools. As such, we named it **DAG Redistribution Lipase 1 (*DRL1*)**, which will be used to denote this gene henceforth. Drl1 contains a GxSxG motif which is conserved among proteins possessing serine hydrolase activity (Figure 2B). Alignment shows that there are 9 proteins in yeast which share this conserved motif (PROSITE PS00120) (Figure 2B). The primary function of known members of the GxSxG containing protein family is lipase activity across a wide variety of substrates. The closest homologs in the yeast genome are Rog1 and Ydl109c at 31.9% and 31.4% identity, respectively (Figure 2B). Although Ydl109c remains uncharacterized, its paralog Rog1 has been shown to have monoacylglycerol activity at the *sn-1* position (Vishnu Varthini et al., 2015). Lpl1 is the nearest evolutionary neighbor to the Drl1 and Rog1/Ydl109c cluster in a phylogenetic tree plotting yeast GxSxG proteins (Figure 2C). Lpl1 is a lipid droplet resident which was shown to exhibit phospholipase B activity on various substrates *in vitro* and it shares 21.1% identity with Drl1 (Selvaraju et al., 2014). Therefore, it is likely that Drl1, as its closest homologs, functions as a type A or B lipase, possibly targeting glycerolipids.

**Figure 2.**
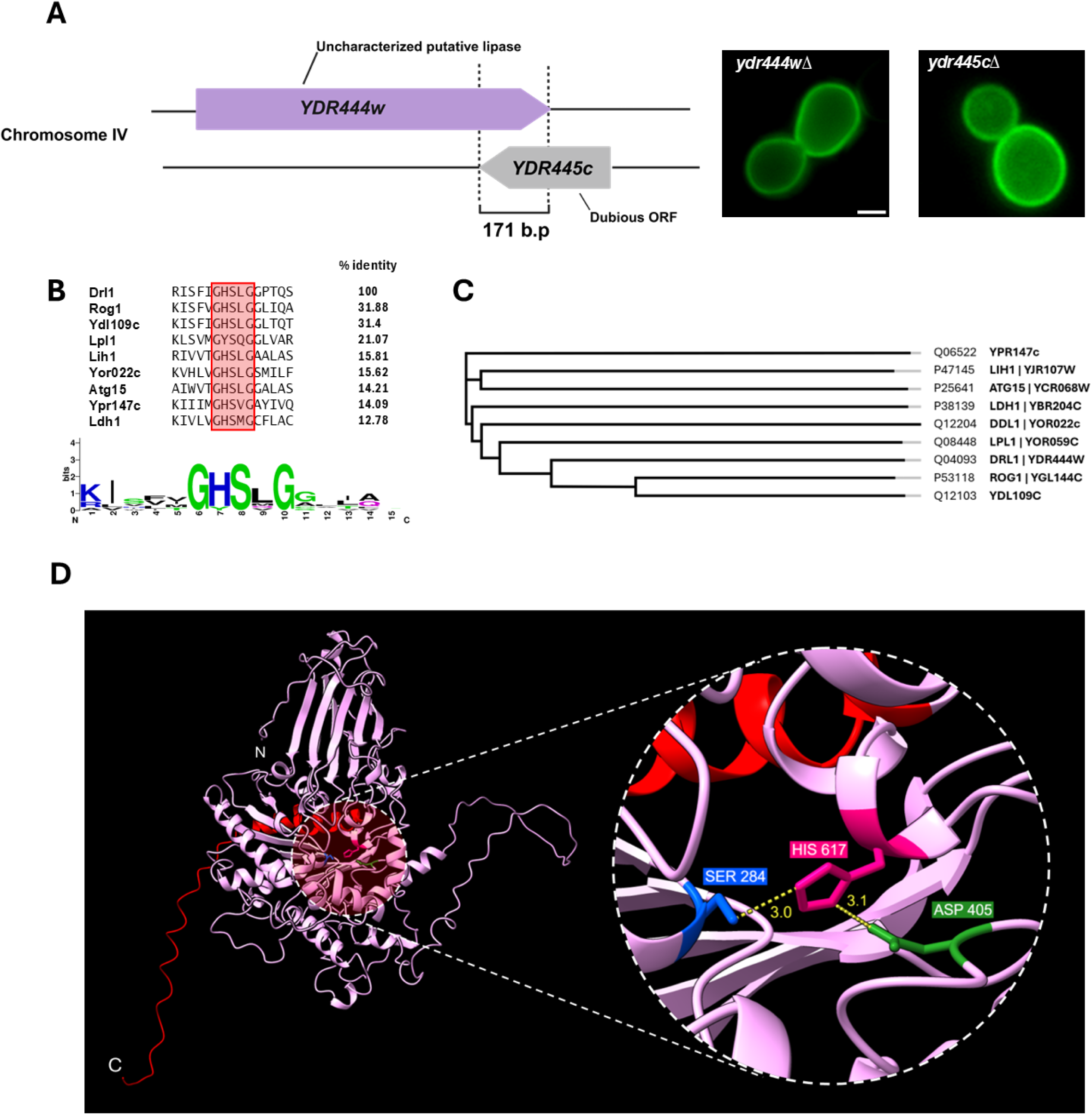
Drl1 is a predicted lipase that regulates cytosolic-facing pools of DAG. **A)** Chromosome arrangement (left) of YDR445c which is annotated as a dubious open reading frame (SGD) and its overlap by 171 b.p with an uncharacterized putative lipase (*YDR444w*). Live microscopy imaging (right) of the fluorescent DAG sensor in *ydr445c*Δ and *ydr444w*Δ mutants grown in defined medium to log phase. Scale bar = 2µm **B)** Sequence alignment of GxSxG family proteins in *S. cerevisiae* including % identity with respect to Drl1. Consensus motif sequence generated using the Weblogo webtool (Crooks et al., 2004) **C)** Phylogenetic tree of GxSxG family yeast proteins indicating evolutionary relationships (Generated in Uniprot Align webtool) **D)** AlphaFold prediction highlighting the canonical serine hydrolase Ser-His-Asp catalytic triad. Serine 284 (blue, within the GxSxG motif), Histidine 617 (pink), and Aspartate 405 (green). C-terminal 57-residue truncation is coloured in red. Distances in Å between sidechains are labeled in yellow.

To assess the structural basis of Drl1 and its similarity to Rog1 and Ydl109c, its predicted AlphaFold 3 (Abramson et al., 2024) structure was analyzed. A canonical serine (Ser) hydrolase typically features a core Ser-Histidine (His)-Aspartate (Asp) catalytic triad, and its mechanism has been described in detail (Rauwerdink & Kazlauskas, 2015). The Ser residue acts as a nucleophile while the proximal His participates as a hydrogen acceptor which is further stabilized by the Asp via hydrogen bonding (Rauwerdink & Kazlauskas, 2015). In the active site of Drl1, this canonical catalytic triad is clearly observed between Ser 284 (within the GxSxG motif), His 617, and Asp 405 (Figure 2D). Measurement of the hydrogen bond participating atoms indicate that all three residues are positioned at a distance conducive for hydrogen bonding (2.7-3.2 Å). Furthermore, this catalytic domain perfectly aligns with that of Rog1 in the predicted 3D structure (Supplementary Figure S2).

In summary, our analysis supports Drl1 as a previously uncharacterized yeast lipase and highlights key predicted structural features essential for its enzymatic activity.

### Changes in Drl1 Abundance or Activity Dramatically Reshape DAG Distribution

We initially identified *DRL1* due to a 171 bp truncation caused by the deletion of the overlapping gene *YDR445C* (Figure 2A). The C-terminal 57-residue region corresponded to a disordered region positioned in the back of the catalytic pocket according to the AlphaFold 3 prediction (Figure 2D). This initial *in silico* analysis guided our decision to generate two mutants of Drl1 to test the functionality of the protein in the context of the observed DAG phenotype. First, to substantiate the contribution of lipase activity to the effect on DAG localization, we mutated G^282^, S^284^, G^286^ within the ^282^GHSLG^286^ serine hydrolase motif to ^282^AHALA^286^ to create a predicted catalytically dead (CD) mutant. Second, ablation of *YDR445C* only accounts for 8.2% of the *DRL1* gene, and we postulated that the C-terminal region was functionally relevant for Drl1. To test this, we introduced a 171 bp (57 aa from G631 to T687) truncation in the carboxy terminus of the protein (Drl1^trunc^). This rationale also guided our decision to avoid fusing tags into the C-terminus of Drl1.

To assess if reversion of the cell periphery DAG phenotype was dependent on Drl1 catalytic activity and C-terminus, we expressed N-terminal V5 tagged Drl1^wt^, Drl1^CD^ and Drl1-^trunc^ proteins from a 2µ plasmid under the constitutive GPD promoter, in *drl1*Δ cells carrying the GFP-DAG sensor. Reintroduction of the WT lipase resulted in a clear shift of the C1δ-GFP probe localizing from the PM to internal compartments of the cell (Figure 3A). Conversely, both the catalytically dead and the truncated mutants were unable to revert the phenotype suggesting that these are both requisites for proper Drl1 function. Western blot confirmed that all three constructs were expressed at similar levels (Figure 3B). Furthermore, integration of a *KanMX-GPD* promoter cassette upstream of *DRL1* was sufficient to cause the DAG-sensor to localize exclusively to internal membrane structures (Figure 3C-D). Localization of the internal DAG signal in these cells was confirmed to colocalize to the vacuolar membrane based on FM4-64 staining (Figure 3E). In contrast with the range of DAG phenotypes displayed by WT cells (with or without KanMX cassette), the localization of the DAG-sensor in *drl1*Δ (cell periphery) and *GPD-DRL1* cells (vacuolar membrane) were extreme and uniformly displayed by all cells (Figure 3D).

**Figure 3.**
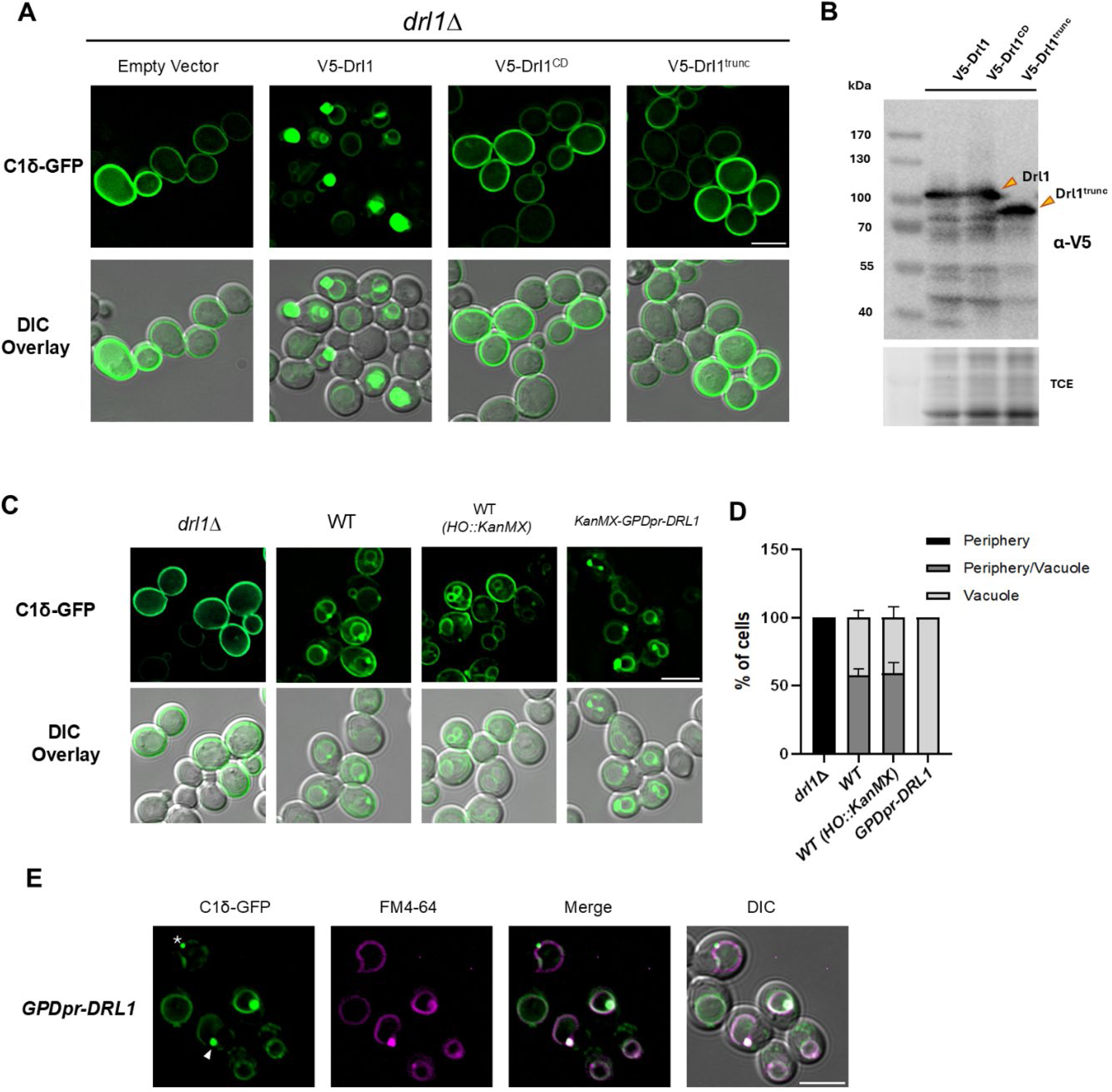
Phenotypic rescue of aberrant DAG localization exhibited by *DRL1* deletion. **A)** Live imaging using fluorescent microscopy of log phase *drl1*Δ cells grown in SD -Leu - Ura expressing WT(V5-Drl1), catalytically dead (V5-Drl1-CD) or truncated (V5-Drl1-*trunc*). DAG is visualized using a C1δ-GFP fluorescent probe. **B)** Cells from the same cultures used for microscopy in A) where lysed and expression of V5-Drl1 WT and mutants was confirmed by Western blot. Proteins were resolved at 140V for 1 hour on a 10% SDS-PAGE gel. Total protein loading was visualized with 2,2,2-trichloroethanol (TCE). Each sample represents 2.6% of the total lysate **C)** Overexpression of Drl1 via GPD promoter swap results in hypermorphic DAG phenotype. Live cell microscopy imaging WT cells (*BY4741*), WT cells containing a kanamycin cassette in the HO locus (*BY4741 HO::KanMX*) and cells overexpressing Drl1 (*KanMX*-*GPDpr-DRL1*). All cells are expressing the C1δ-GFP DAG sensor (grown in SD -Ura). Scale bar = 5 µm **D)** Quantification of DAG phenotype classified as either having C1δ-GFP signal exclusively in the vacuole, the cell periphery or in both the cell periphery and vacuole. Bars represent ± SD for three independent experiments. **E)** DAG sensor (C1δ-GFP) co-localization with vacuolar dye (FM4-64) in cells overexpressing Drl1 by swapping the native *DRL1* promoter for the constitutive *GPD* promoter. Log phase cells grown in SD -Ura were stained at 30 °C for 45 minutes with 32 μM FM4-64 and imaged immediately. Asterisk denotes DAG vacuolar microdomain. Arrowhead points to a prevacuolar compartment enriched in DAG. Scale bar = 5 µm.

Overall, our findings underscore the essential role of Drl1 in maintaining proper DAG distribution and confirm that both the catalytic GxSxG motif and the disordered C-terminus are critical for its enzymatic activity.

### Drl1 localizes to nucleus and plasma membrane

To further characterize Drl1, we next interrogated its localization by inspecting *drl1*Δ cells expressing GFP-Drl1 or GFP-Drl1^CD^ actively growing in defined medium containing 2% glucose (Figure 4). High-resolution images captured GFP-Drl1 localized to the PM and concentrated in the nucleus as shown by co-localization of GFP-Drl1 with the nucleolar marker Sik1-RFP (Figure 4 A&B). Interestingly, cells expressing the catalytically dead version Drl1^CD^ displayed bright foci and showed decreased nuclear signal and lack of PM localization (Figure 4A). Expression levels of Drl1 and Drl1^CD^ were comparable, and subcellular fractionation confirmed that both proteins are partially associated with the insoluble/membrane fraction (Figure 4C).

**Figure 4.**
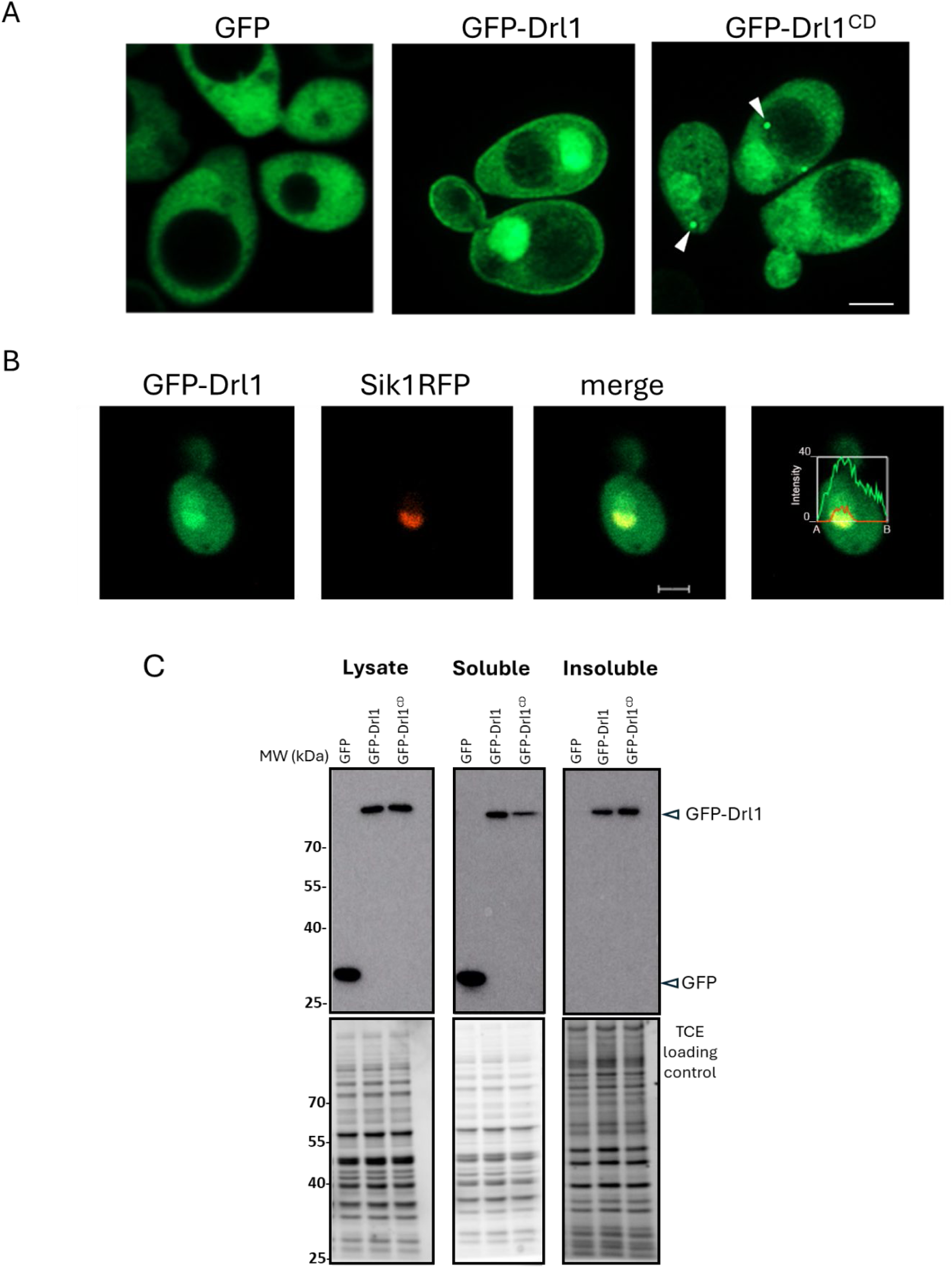
Drl1 localizes to nucleus and plasma membrane. **A)** Airyscan high-resolution microscopy from *drl1*Δ cells transformed with pRS416-GPDpr-GFP; pRS416-GPDpr-GFP-*DRL1* or pRS416-GPDpr-GFP-*DRL1*^CD^. Arrowheads point to GFP-*DRL1*^CD^ foci. **B)** Confocal microscopy showing WT cells expressing Sik1-RFP from its endogenous locus transformed with pRS416-GPDpr-GFP-*DRL1*. The red and green lines in the graph show the intensity of the RFP and GFP signal along the line indicated from A to B, respectively. Scale bar: 2 µm. **C)** Subcellular fractionation of *drl1*Δ cells transformed as in A) was performed as indicated in Materials & Methods. Equal amounts of proteins from each lysate, soluble and insoluble fractions per sample were separated by SDS-PAGE and western blotted using an α-GFP monoclonal antibody. Total proteins (loading control) were visualized with 2,2,2-trichloroethanol (TCE).

This dual PM-nuclear localization of Drl1 is unique among yeast GxSxG containing proteins (Figure 2B), none of which emerged as hits in our screen, suggesting that Drl1’s specific localization, and thus its territory of action, is critical for proper DAG distribution.

### Drl1 impacts sterol distribution

Based on the observation that the cell periphery category contained genes involved in PS, phosphoinositides (PIPs), and sterol metabolism as hits, we decided to investigate if lack of Drl1 would also influence their localization. For this, a collection of lipid binding probes, validated for use in *S. cerevisiae,* were monitored in cells lacking or overexpressing Drl1. No differences in probe distribution were observed between all three strains expressing Spo20-GFP (phosphatidate, PA) (Nakanishi et al., 2004); Lact-C2 (PS) (Yeung et al., 2008); Fapp1-PH-GFP (PI4P) (Stefan et al., 2002) and PLC-PHδ-GFP (PI4,5P_2_) (Stefan et al., 2002) (Figure S3). By contrast, sterol localization was markedly altered. As previously shown, WT cells exhibited a strong polarized signal of the sterol probe GFP-ALOD4 at the PM of daughter cells (Chauhan & Fairn, 2021; Sosa Ponce et al., 2025). In *drl1*Δ cells, we observed an approximately eight-fold increase in the percentage of cells displaying a soluble ALOD4 phenotype in comparison to the WT (p < 0.01) (Figure 5). Overexpression of Drl1 (GPDpr*-DRL1*) partially restored sterol localization to WT yeast, with a small but significant increase in puncta and cytosol signal, at 20% and 15%, respectively (Figure 5).

**Figure 5.**
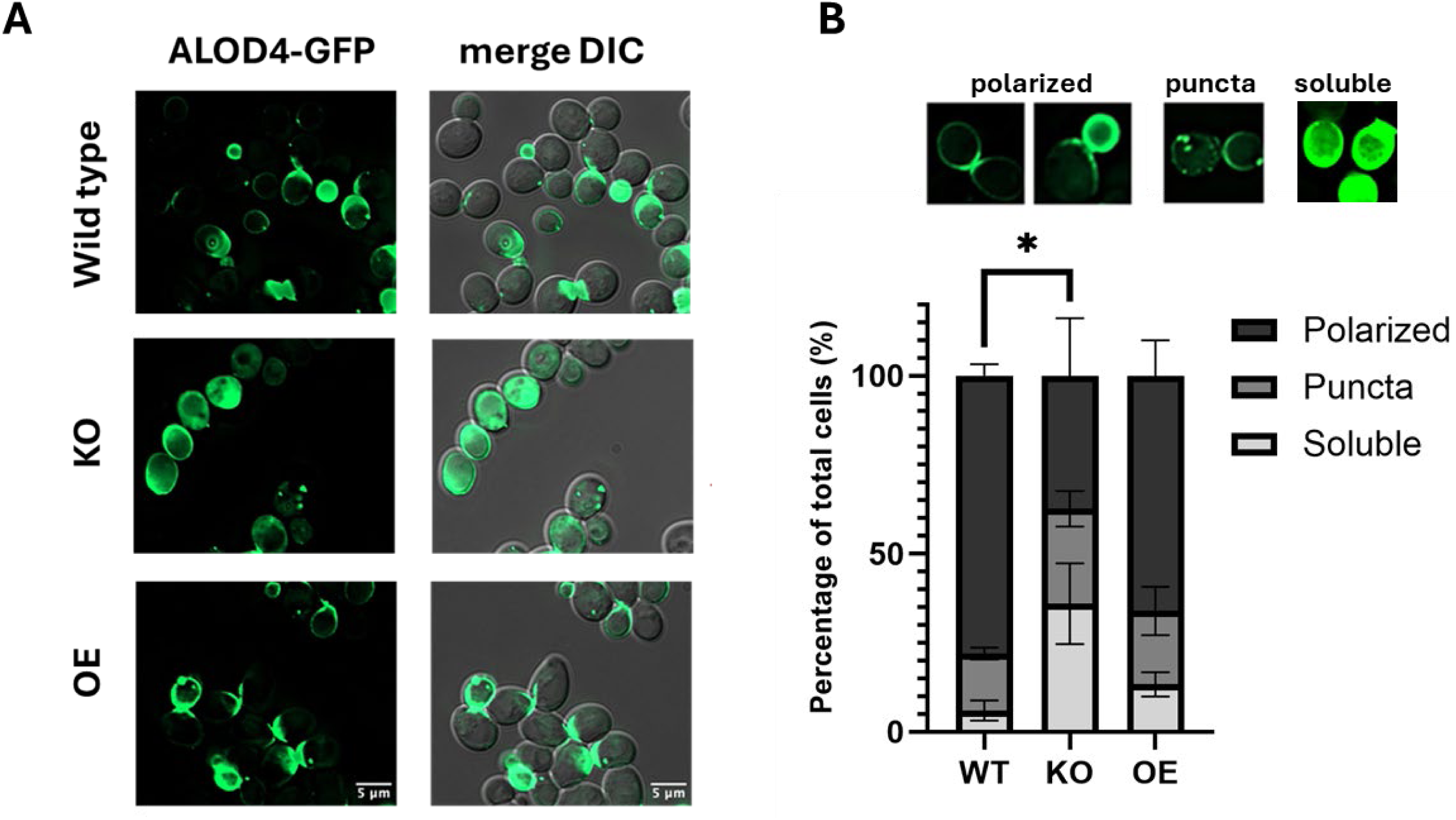
Drl1 influences sterol localization. **A)**. Wild-type (WT), *drl1*Δ (KO) and *DRL1* overexpressing (OE) cells were transformed with a GFP-ALOD4 expression plasmid to monitor cytosolic facing pools of sterols. Transformants were grown to mid-log phase in SD -Ura followed by 30 minutes SD -Ura -Met to induce GFP-ALOD4 expression. Scale bar = 5 μm. **B)** Quantification of images from A, based on the indicated categories observed in cells expressing the GFP-ALOD4 sterol probe: puncta, soluble, and polarized expressed as percentage of total cells. Bars represent average ± SD of 3 experimental replicates, each counting 50 cells (* = p< 0.05).

These results indicate that altered levels of Drl1 specifically affect sterol polarization while it has no significant effect on PA, PS, PI4P and PI4,5P_2_ when actively growing in complete defined medium.

### Drl1 modulates lysolipid and unsaturated fatty acid levels

To investigate the role of Drl1 in lipid metabolism and homeostasis we next performed untargeted lipidomics in *DRL1* knockout (*Δdrl1::KanMX*) and *DRL1* overexpressing (*KanMX-GPDpr-DRL1*) strains. An isogenic WT strain carrying an integrated KanMX cassette in the HO locus was used as control (*HO::KanMX*).

Lipidomic analysis in triplicates was performed (Figure 6A and Supplementary Table S2) and PCA analysis reflected distinct clustering for each set of replicates, indicating that lipidomic variation among each experimental condition was distinct (Figure 6B). WT samples were flanked by the knockout and overexpression conditions. Volcano plots between *GPDpr*-*DRL1* (OE) and *drl1*Δ (KO) were analyzed to identify lipid species based on fold change and statistical significance (Figure 6C). Cutoffs for a lipid to be significant were a fold change greater than 1.5 and a p value less than 0.05.

**Figure 6.**
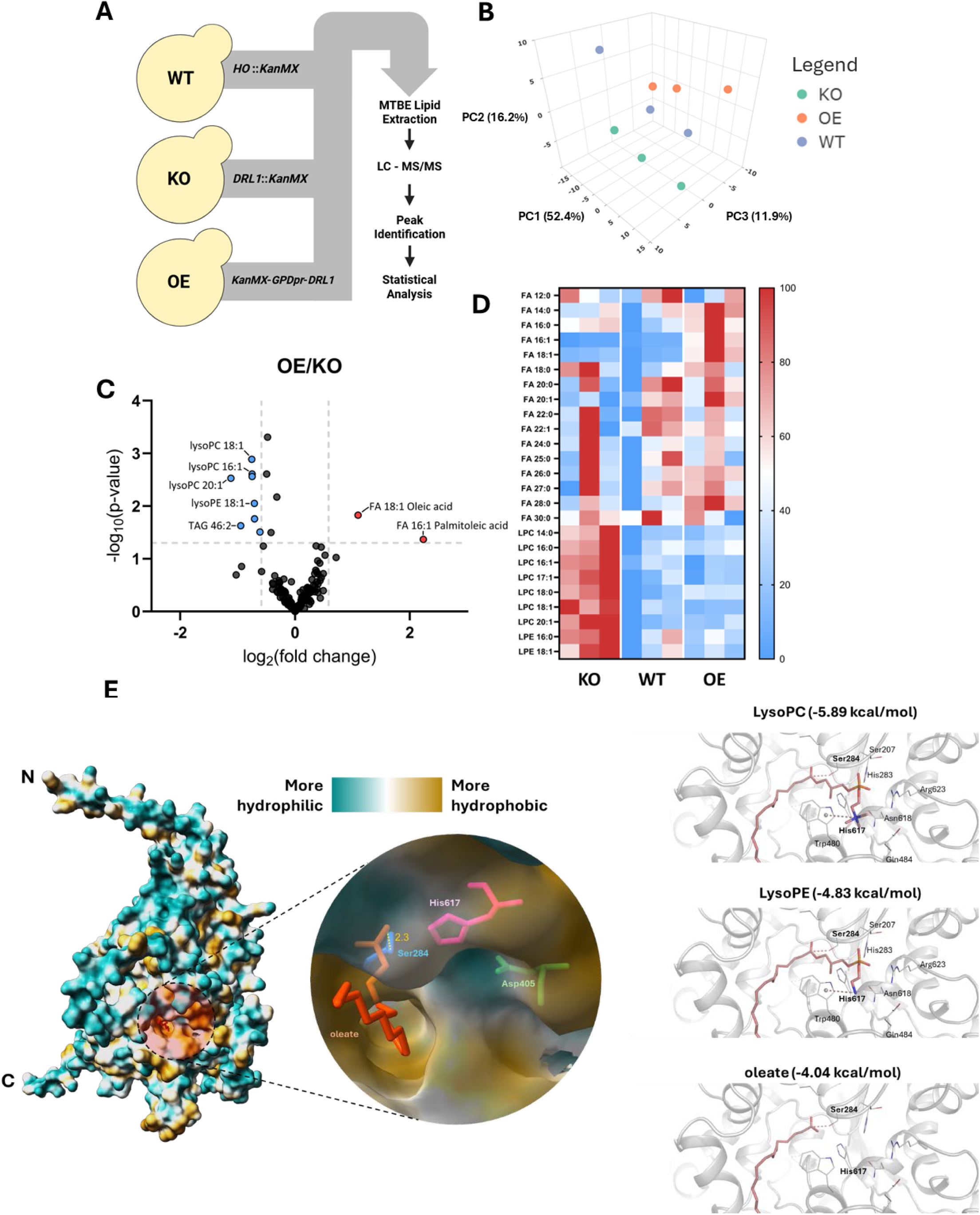
Drl1 modulates fatty acid and lysolipid levels. **A)** Flowchart of experimental conditions and lipidomics pipeline. **B)** Principal component analysis of lipidomic profiles of *DRL1::KanMX* cells (KO), *KanMX*-*GPDpr-DRL1* overexpression cells (OE) or WT *HO::KanMX* cells (WT). **C)** Volcano plots comparing lipid species abundance between OE and KO conditions, plotting −log10(p-value) versus log2(fold change). Lipids significantly decreased or increased in OE relative to KO are shown in blue and red, respectively (Cutoffs: fold change > 1.5, p-val < 0.05). **D)** Heat map of fatty acid (FA), lysophosphatidylcholine (LPC), and lysophosphatidylethanolamine (LPE) species detected across WT, KO, and OE conditions. Abundance values were normalized per lipid species by setting minimum and maximum values to 0 and 100, respectively, with intermediate values expressed as percentages.**E) Left**: AlphaFold 3 prediction of Drl1 modeled with oleate. Structure is coloured by surface hydrophobicity (colouring key is shown). The canonical serine hydrolase Ser-His-Asp catalytic triad is shown, with Serine 284 (blue, within the GxSxG motif), Histidine 617 (pink), and Aspartate 405 (green). Oleate is shown in orange. Distance in Å between oleate and Ser284 is labeled in yellow. **E) Right**: Molecular docking of LysoPC, LysoPE, and oleate into Drl1, showing the predicted binding affinity expressed as the estimated free energy of binding (in parentheses). Residues Ser284 and His617 from the catalytic Ser–His–Asp triad are indicated. Additional residues predicted to interact with each lipid headgroup within 5 Å are shown. Dashed lines represent predicted polar interactions.

The first observation was that the null mutant (*drl1*Δ) accumulated Lyso-PC and Lyso-PE species with unsaturated acyl tails of varying lengths (Figure 6C). The most significant enriched lipids when comparing KO to OE were Lyso-PC (18:1) (p-value = 0.0025) and Lyso-PC (16:1) (p-value = 0.0013) (Figure 6C and S4). Interestingly, overexpression of Drl1 resulted in elevated fatty acids (FAs), namely oleic acid (18:1) and palmitoleic acid (16:1) (Figure 6C and S4). Plotting all FA’s and lysolipids on a heatmap further illustrates the point that all LysoPC and LysoPE species, regardless of acyl chain lengths and degree of saturation, are increasing in abundance when *DRL1* is deleted (Figure 6D). This is not the case for FA’s where a pronounced increase of the unsaturated fatty acids (16:1 & 18:1) stands out (Figure 6D). Based on these results, the most obvious hypothesis is that Drl1 is a lysophospholipase which acts on monounsaturated LysoPC/LysoPE releasing monounsaturated fatty acids. Because attempts to detect LysoPC/LysoPE lipase activity using recombinant Drl1 or membrane extracts from Drl1 overexpressing yeast have so far been unsuccessful, we turned to structural modeling with the AlphaFold 3 deep learning algorithm to explore potential Drl1:lipid interactions *in silico.* Remarkably, oleate (18:1), which was available for modelling in this platform, was predicted to interact with Ser284 of the consensus pentapeptide GxSxG and with His617 from the Ser-His-Asp catalytic triad within a hydrophobic pocket of Drl1 containing the catalytic domain (Figure 6E). While AlphaFold3 has greatly improved our ability to predict biomolecular complex structures, the model cannot predict binding affinity, a key property underlying molecular function. Therefore, we next turned to perform molecular docking, using Boltz-2, an open-source deep learning framework for both biomolecular structure and affinity prediction (Passaro et al., 2025). In addition to oleate (18:1) we also included LysoPC (18:1) and LysoPE (18:1) for estimating Drl1-lipid binding affinity. Boltz-2 utilizes the performance of free-energy perturbation methods. The obtained predicted binding free energies support the hypothesis that Drl1 preferentially recognizes lysophospholipids, consistent with a LysoPC/LysoPE lipase–like function. LysoPC shows the strongest binding, followed by LysoPE, while oleate binds substantially more weakly (LysoPC > LysoPE > oleate). The enhanced affinity for LysoPC and LysoPE is suggested to be driven by favorable van der Waals interactions, hydrogen bonding, and desolvation, along with headgroup-specific interactions involving conserved active-site residues such as Ser284 (Figure 6E-left). In contrast, oleate centered predictions lack these stabilizing contacts. It is worth noting that bulkier lipids such as DAG are sterically excluded from the Drl1 binding pocket. Together, these results indicate that Drl1 is selectively tuned for lysophospholipid substrates, supporting its proposed role as a LysoPC/LysoPE-specific lipase, although this proposed activity remains to be experimentally validated.

### LysoPC is a counter spatial regulator of DAG and sterols

The findings from the lipidomics analysis raised the question as to how lack of Drl1 results in DAG and sterol distributions deviating from WT. Since lysolipids (LysoPC and LysoPE) accumulate in *drl1*Δ, we hypothesized that they may alter the lipid environment in a manner that affects the spatial distribution of DAG and sterols. Indeed, consistent with our observations in *drl1*Δ cells, we previously reported that treatment of WT yeast with non-metabolizable LysoPC analogues led to loss of spatial polarization and reduction of sterols at the plasma membrane (Czyz et al., 2013; Sosa Ponce et al., 2025; Zaremberg et al., 2005). We wondered if these metabolically stable LysoPC analogues would also alter DAG distribution as displayed by cells lacking Drl1. We therefore next assessed the localization of the DAG sensor in WT cells treated with the lysoPC analogue edelfosine (1-O-octadecyl-2-O-methyl-rac-glycero-3-phosphocholine). This LysoPC analogue is metabolically stable due to the presence of an ether linkage in the *sn-1* position preventing its de-acylation as well as a methoxy group in the *sn-2* position which cannot be acylated (Figure 7A). These modifications lead to edelfosine accumulation in yeast allowing us to imitate the increase of LysoPC observed in *drl1*Δ cells. Thus, we next assessed the effect of edelfosine on DAG spatial distribution in WT yeast (Figure 7A). Remarkably, edelfosine treatment mimicked the *drl1*Δ phenotype inducing a radical shift in the DAG probe localization, moving from the vacuolar pool to the cell periphery (Figure 7A).

**Figure 7.**
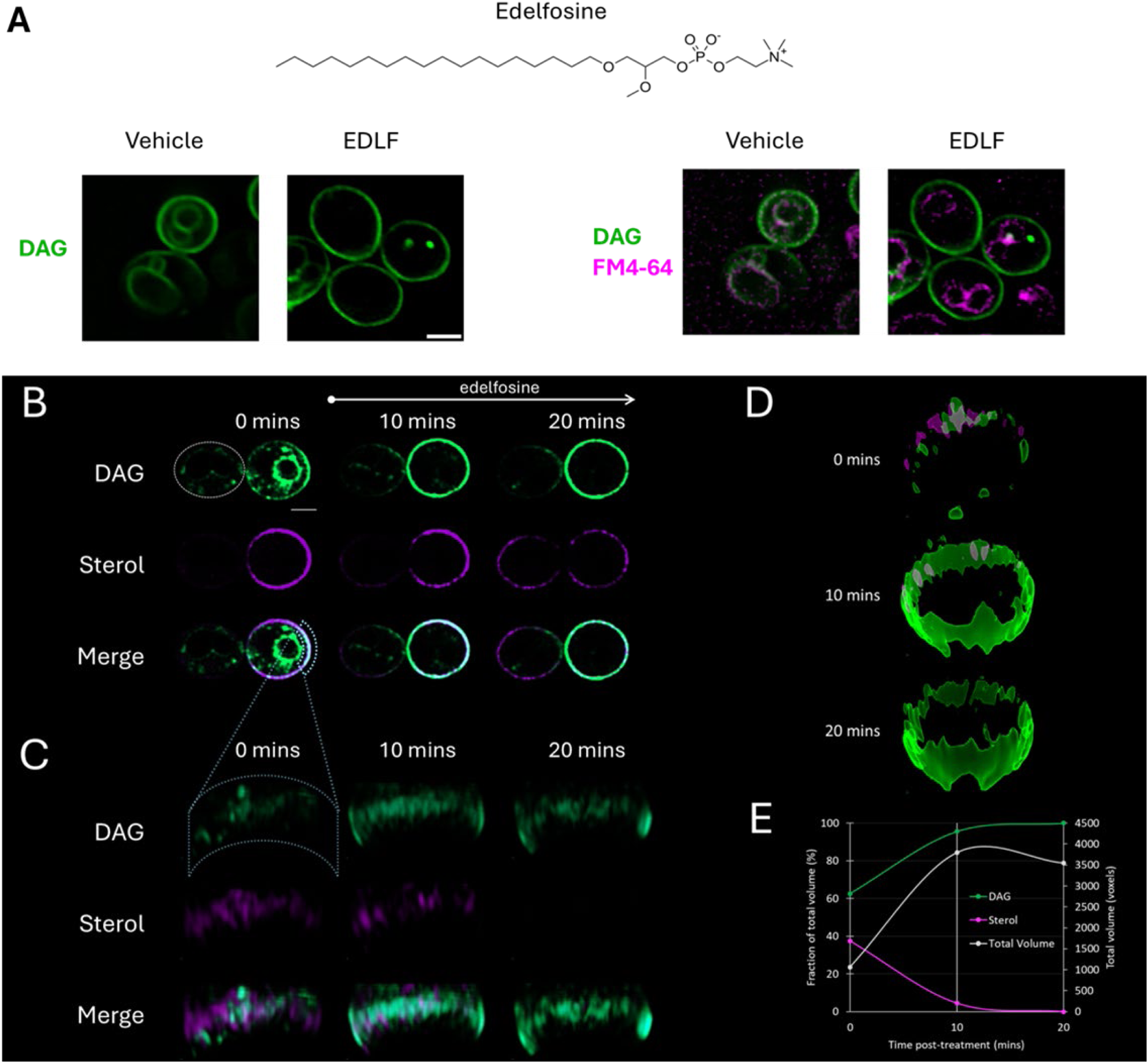
Non-metabolizable LysoPC analogue alters DAG and sterol distribution at the PM. **A)** Structure of Edelfosine, a non-metabolizable LysoPC analogue. For Edelfosine treatment (EDLF), 20 μM of the drug was added to WT cells expressing the DAG probe (C1δ-GFP) and preloaded with FM4-64 to visualize vacuoles. Cells were incubated at 30 °C for 20 minutes before imaging with epifluorescence microscope. Scale bar= 2 um. **B)** Representative Airyscan images of a WT (BY4741) budding cell expressing the DAG probe, C1δ-GFP, and the sterol probe, mCherry-ALOD4 from plasmids. Cells were actively growing in defined selective SD-ura-leu medium while imaged. The times post-edelfosine (non-metabolizable LysoPC analogue) addition to the media are indicated. Top dashed circle (white) denotes mother cell periphery. Bottom dashed format (light blue) indicates the plasma membrane (PM) area selected for 3D analysis in panel B. Scale bar= 2 µm. **C)** 3D rendering of chosen PM areas (at lower exposure than panel A to avoid saturation), with 15 slices representing 2.66 μm thickness. **D)** Objects representing the volume (voxels) of the entire bud PM, occupied by the DAG (green) and sterol (magenta) domains over time. **E)** Quantification of lipid domain volumes expressed as ratio of individual lipid volumes to total volume over time of treatment with edelfosine, as outlined in Materials and Methods.

To obtain more details of the effect of edelfosine on both sterol and DAG pools, WT cells simultaneously expressing mCherry-ALOD4 and C1δ-GFP probes were imaged during a short period of edelfosine treatment. High-resolution live imaging revealed that DAG and sterol probes exhibited polarized localization at the bud PM at time zero, forming a patchy distribution with distinct microdomains for both lipids (Figure 7B). Strikingly, a brief 10-minute treatment with edelfosine caused the DAG sensor to diminish in the vacuole and become even more enriched at the PM (Figure 7C). These changes were accompanied by a concomitant reduction of the sterol probe at the bud PM, consistent with previous observations in edelfosine-treated cells (Czyz et al., 2013; Sosa Ponce et al., 2025; Zaremberg et al., 2005). A 3D rendering of a region of the bud PM revealed an intricate, interdigitated arrangement of unique DAG and sterol-enriched territories at time zero. Sterol domains dissipated upon edelfosine treatment while DAG presence expanded (Figure 7D). Volume quantification of the entire bud PM showed that, initially, it contained roughly similar volumes of DAG and sterols (Figure 7E). By 10 minutes with edelfosine, sterol volume had rapidly declined to zero, while DAG volume increased. Notably, the total DAG plus sterol lipid-occupied volume expanded more than threefold between 0 and 10 minutes (Figure 7E). These findings indicate that DAG not only replaces the volume vacated by sterols but also exceeds the original combined volume of both lipids.

Altogether, our results support a role for LysoPC as a spatial regulator of DAG, also affecting the PM’s sterol retention capacity.

## Discussion

Eukaryotic cells rely on remarkable versatility in their membrane lipid composition to maintain homeostasis and support growth in the face of changing environmental and metabolic demands. This flexibility allows membranes to preserve optimal fluidity, curvature, permeability, and signaling capability, ensuring that essential processes proceed efficiently.

In this study, we show that cytosolic-facing DAG pools in yeast contribute to this functional versatility by providing a mobile lipid source that can be rapidly deployed to counteract phospholipid imbalance in cellular membranes.

To uncover regulators of DAG spatial organization, we performed a visual genome-wide screen to identify perturbations in yeast DAG dynamics. This approach revealed a discrete number of mutants with abnormal DAG localization. Among them, a mutant lacking an uncharacterized putative lipase, which we named Drl1, emerged as a notable hit, displaying a striking phenotype in which DAG remained persistently localized at the cell periphery. In addition, sterols were displaced in the *drl1*Δ mutant. Biochemical characterization of this mutant ultimately revealed that LysoPC functions as a key spatial regulator of cytosolic-facing DAG pools. We show that LysoPC accumulates in *drl1*Δ cells and that a non-metabolizable analog of LysoPC triggers a directed redistribution of these DAG pools in WT yeast, a process that is tightly synchronized with the retrograde mobilization of sterols from the PM. Our findings support a model in which LysoPC perturbs local PM packing, driving sterol efflux and, in turn, promoting the flow of DAG from the vacuolar pool.

A rapidly diffusing pool of sterols is increasingly recognized as a defining feature of PM lipid organization in living cells, highlighting the dynamic nature of sterol behavior within bilayers (Doktorova et al., 2025). Across eukaryotes, from yeast to mammals, the pronounced sensitivity of PM sterol asymmetry to lipid imbalance underscores how tightly sterol distribution is coupled to the broader phospholipid environment (Doktorova et al., 2025; Mondal et al., 2008; Solanko et al., 2018). For example, perturbations in sphingolipids (SL) are well known to trigger sterol relocalization away from the PM (Das et al., 2014; Doktorova et al., 2025; Mondal et al., 2008). Under similar conditions of SL imbalance, DAG has been proposed to accumulate in the cytosolic leaflet of the PM, where it can activate PKC signaling in mammalian cells (Ueda et al., 2013).

Sterol influx is known to be triggered by LysoPC analogues such as edelfosine, platelet activating factor (PAF) (Sosa Ponce et al., 2025; Zaremberg et al., 2005; Zaremberg & McMaster, 2002), and other small amphipathic molecules (Tettamanti et al., 2025). Although the mechanism by which LysoPC induces sterol influx is not fully understood, it has been shown that lysolipids can form hydrogen bonds with sterols (Krasnobaev et al., 2022) likely altering sterol interactions with other PM lipids.

Because sterol depletion compromises PM packing and function, cells may compensate by DAG translocation to the PM, leveraging its conical shape and relative hydrophobicity as a functional surrogate. This compensatory DAG influx aligns with the broader principle that sterols and DAG are non-bilayer forming lipids which modulate membrane curvature and packing (Yu et al., 2021). However, DAG has a weaker membrane condensing capacity, which means that higher concentrations are required to achieve ordering effects comparable to sterols (Alwarawrah et al., 2012).

Taken together, DAG appears to be a rapid response mechanism for dealing with perturbations in membrane architecture induced by LysoPC. This lipid co-movement provides a dynamic conduit for rapidly reconfiguring membrane composition in response to disrupted lipid homeostasis.

Sterol retrograde transport induced by small amphipathic molecules has recently been linked to Lam2/4-dependent non-vesicular transport at PM-ER contact sites, which operates significantly faster than conventional membrane trafficking routes (Tettamanti et al., 2025). In contrast, our yeast genome-wide survey indicates that DAG distribution relies on an interconnected network involving components of the CVT, MVB, and ESCRT pathways rather than non-vesicular transport mechanisms. These pathways converge at the vacuole and share key machinery like the ESCRT complexes, which mediate membrane-remodeling events in which membranes bud away from the cytosol (Hurley & Hanson, 2010). Yeast studies have shown that ESCRT-III proteins are essential for maintaining normal PS and PE levels, and mutants defective in ESCRT-III exhibit altered lipid pools, including reduced PS and PE and elevated DAG (Jorgensen et al., 2020). Together, these observations lead us to propose that although both sterols and DAG can rapidly flip-flop between membranes (Bennett & Tieleman, 2012), lysolipid imbalance at the PM reduces its sterol-retention capacity, triggering rapid non-vesicular sterol retrograde transport and DAG redistribution via anterograde vesicle-mediated trafficking from the vacuole to the PM.

It is striking that the *drl1*Δ mutant maintains normal growth despite elevated LysoPC/LysoPE levels and pronounced defects in DAG and sterol distribution-conditions that may compromise membrane integrity and signaling if persistent in time. This resilience suggests that the mutant engages compensatory lipid-homeostatic pathways. A negative genetic interaction has been reported between *drl1*Δ and *psd1*Δ, which lacks the mitochondrial PS decarboxylase, implying that Drl1 may contribute to the pathway that activates the vacuolar/Golgi PS decarboxylase Psd2 when ethanolamine is limiting (Y. Wang et al., 2020). In line with this possibility, our visual genetic screen identified both, *PSD1* and *PSD2,* as modulators of DAG distribution: *psd1*Δ phenocopied Drl1 overexpression, whereas *psd2*Δ mirrored the *drl1*Δ phenotype. The differential contribution of Psd1 and Psd2 to DAG spatial distribution, as well as their relationship with Drl1, is intriguing and warrants further investigation to fully understand the underlying mechanisms.

Loss of *DRL1* (*drl1*Δ) and *DRL1* overexpression led to the accumulation of LysoPC/LysoPE and unsaturated fatty acids, respectively. These phenotypes suggest that Drl1 contributes to lysolipid turnover, a process that is highly redundant in yeast and largely mediated by *PLB1/2/3* and *NTE1*. In addition, nine GxSxG domain containing proteins in yeast are type A and B lipases (Debelyy et al., 2011; Manda et al., 2018; Selvaraju et al., 2014; Urafuji & Arioka, 2016; Vishnu Varthini et al., 2015). Notably, none of these lipases emerged as hits in our screen, suggesting that Drl1’s unique dual localization at the PM and nucleus may position it to act on lysolipid pools that regulate DAG distribution.

In summary, our findings have broad implications for eukaryotic cells, from yeast to mammals, as the coordinated movement of DAG and sterol pools potentially reflects a conserved principle of membrane homeostasis. The demonstration that LysoPC can trigger DAG-sterol counter trafficking suggests that this is a regulatory mechanism that enables cells to sense and correct phospholipid imbalance. More globally, our work contributes to the emerging interconnectivity map of various lipids and their counter-balancing capacity to preserve PM homeostasis.

## Data Availability

Data reported in this article are available in the published article and its online supplemental material. Strains and plasmids are available from the corresponding author upon request.

## Acknowledgements

We wish to thank Dr Adriana Zardini Buzatto for her guidance in the analysis of lipidomics data and Claire Mallard for her technical contributions related to Supplementary Figure S3. We would also like to thank Aaron Neiman, Scott Emr and Greg Fairn for their kind gifts of plasmids. We thank Lihi Gal for her help with the yeast high content screen. We thank Dr. Maxim Itkin and Dr. Sergey Malitsky from the Weizmann Institute of Science Metabolomics unit for performing the lipidomic analysis.

This work was supported by a Natural Sciences and Engineering Research Council of Canada Discovery Grant (NSERC 2023-04320) and Alberta Innovates (AI 242506270) to VZ, and a CIHR Grant (MOP-142403) to KB. PP was supported by a grant from the University of Buenos Aires (UBACYT 20020190100122BA).

This study was supported by a Chan Zuckerberg Initiative (CZI) grant (2023-331952) as well as the Institute for Environmental Sustainability (IES) of the Weizmann Institute of Science. The robotic system of the Schuldiner lab was purchased through the kind support of the Blythe Brenden-Mann Foundation. MS is an Incumbent of the Dr. Gilbert Omenn and Martha Darling Professorial Chair in Molecular Genetics.

## Author Contributions

Conceptualization: *S. Ganesan, V. Zaremberg and M Schuldiner;* Formal Analysis*: A. Henderson, A. Lalani, S. Ganesan, N.Zung, P. Portela, H. Mesa-Galloso V. Zaremberg and M. Schuldiner;* Funding Acquisition*: K. Baetz; P. Portela, V. Zaremberg and M. Schuldiner;* Investigation: *A. Henderson, A. Lalani, S. Ganesan, M.L. Sosa Ponce, N.Zung, P. Portela, H. Mesa-Galloso, V. Zaremberg and M Schuldiner;* Methodology: *A. Henderson, A. Lalani, S. Ganesan, M.L. Sosa Ponce, N.Zung, P. Portela, H. Mesa-Galloso, V. Zaremberg and M Schuldiner;* Project Administration*: V. Zaremberg and M Schuldiner;* Resources: *K. Baetz; P. Portela, V. Zaremberg and M. Schuldiner;* Supervision: *S. Ganesan, M.L. Sosa Ponce, V. Zaremberg and M. Schuldiner;* Validation: *A. Henderson, A. Lalani, S. Ganesan;* Visualization: *A. Henderson, A. Lalani, S. Ganesan, V. Zaremberg and M. Schuldiner;* Writing-original draft: *A. Henderson and V. Zaremberg;* Writing-review and editing*: all authors*.

## Materials and Methods

### Reagents

Unless stated otherwise, all reagents were purchased from Fisher or Sigma-Aldrich. Yeast extract peptone, and yeast nitrogen base were sourced from MP biomedicals. Edelfosine was a kind gift from Medmark Pharma GmbH.

### Yeast, strains, plasmids and primers

Detailed information of yeast strains, plasmids, and primers used in this study is provided in Tables 1-3. A catalytically dead mutant of *DRL1* was constructed by introducing four sequential point mutations (G845C, T850G, A858C, G857C) into a pDONR221-*DRL1* entry clone (DNASU: ScCD00008940) using the QuikChange Lightning Multi Site-Directed Mutagenesis Kit (Agilent). A 171 b.p, C-terminal truncation was introduced by PCR amplifying *DRL1* with gateway compatible primers (Gtwy-*YDR444w-*F, Gtwy-*YDR444w*Trunc-R). The amplified product was then cloned into a Gateway pDONR221 donor vector (ThermoFisher Scientific) to create an entry clone. Both the catalytically dead and truncated entry clones were confirmed by Sanger sequencing (DNA sequencing facility, University of Calgary). Entry clones were then subcloned into the gateway compatible destination vector pZM552 (DNASU: EvNO00335423). The *S. cerevisiae* Advanced Gateway Destination Vectors (Alberti et al., 2007) were kindly gifted by Susan Lindquist (kit #1000000011; Addgene). The native promoter of *DRL1* was swapped using the PYM-N14 plasmid (Janke et al., 2004) as template DNA for PCR amplification of the KanMX-GPD cassette using S1 (*YDR444w*-GPD-SWAP-S1-F) and S4 (*YDR444w*-GPD-SWAP-S4-R) site specific primers (Table 2.3). Integration was confirmed by colony PCR using a forward primer specific to the kanamycin cassette (*KanMX4*-ATG+26-F) and a reverse primer specific to the *DRL1* gene locus (*YDR444w*-1440-R) (Table 2.3).

**Table 1.** Strains used in this study.

| Number | Name | Genotype | Mating Type | Background | Source |
| --- | --- | --- | --- | --- | --- |
| VZY125 | KanMX-GPDpr-DRL1 | *YDR444w::KanMX-GPDpr-YDR444w his3Δ1 leu2Δ0 met15Δ0 ura3Δ0* | MAT a | BY4741 | This study |
|  | BY4741 | *his3Δ1 leu2Δ0 met15Δ0 ura3Δ0* | MAT a | BY4741 | (Brachmann et al., 1998) |
|  | BY4741 (*HO::KanMX*) | *his3Δ1 leu2Δ0 met15Δ0 ura3Δ0 HO::KanMX* | MAT a | BY4741 | Schuldiner's lab |
|  | *drl1Δ* | *his3Δ1 leu2Δ0 met15Δ0 ura3Δ0 YDR444w::KanMX* | MAT a | BY4741 | (Giaever et al., 2002) |
|  | SGA Query strain | *his3Δ1 leu2Δ0 met15Δ0 ura3Δ0 can1Δ::STE2pr-spHIS5 lyp1Δ::STE3pr-LEU2 Vam6-mCherry::NatR* | MAT α |  | Schuldiner's lab |
|  | Yeast DAMP (Decreased Abundance by mRNA Perturbation) library |  |  |  | Breslow et al., 2008 |
|  | Yeast deletion library |  |  |  | (Giaever et al., 2002) |

**Table 2.** Plasmids used in this study.

| Name | Description | Source |
| --- | --- | --- |
| pGPD416-GFP-C1-PKCδ | *GPD* promoter, GFP-C1- PKCδ (CEN, *URA3*) | (Ganesan et al., 2016) |
| pGPD415-mCherry-ALOD4 | *GPD* promoter, mCherry-ALOD4 (CEN, *LEU2*) | This study |
| pUG36-GFP-ALOD4 | *MET25* promoter, GFP-ALOD4 (CEN, *URA3*) | (Chauhan & Fairn, 2021) |
| pGPD416-GFP-Lact-C2 | *GPD* promoter, GFP-Lact-C2 (CEN, *URA3*) | (Yeung et al., 2008) |
| pRS426-G20 | *TEF2* promoter, GFP-Spo20<sup>51-91</sup> (2 μ, *URA3*) | (Nakanishi et al., 2004) |
| pRS425-GFP-PH-FAPP | *PRC1* promoter, GFP-PH-FAPP1 (2 μ, *LEU2*) | (Stefan et al., 2002) |
| pRS426-GFP-2xPH-PLCd | *PRC1* promoter, GFP-2xPH-PLCd (2 μ, *URA3*) | (Stefan et al., 2002) |
| pRS416-GPD-GFP-*ccdB* | Destination Vector: N-tagged GFP (CEN, *URA*) | (Alberti et al., 2007) |
| pRS416-GPD-GFP-DRL1 | *GPD* promoter, N-tagged GFP *DRL1* (CEN, *URA*) | This Study |
| pRS416-GPD-GFP-DRL1(CD) | *GPD* promoter, N-tagged GFP *DRL1* CD (CEN, *URA*) | (Alberti et al., 2007) |
| pDONR221-DRL1 | Entry clone, *DRL1* | DNASU: ScCD00008940 |
| pZM425-GPD-3xV5-*ccdB* | Destination Vector, N-tagged 3xV5, (2μ, *LEU*) | DNASU: EvNO00335423 |
| pZM425-GPD-3xV5-DRL1 | *GPD* promoter, N-tagged 3xV5 *DRL1* (2μ, *LEU*) | This Study |
| **pZM425-GPD-3xV5-DRL1 (CD)** | *GPD* promoter, N-tagged 3xV5 *DRL1* CD (2μ, *LEU*) | This Study |
| **pZM425-GPD-3xV5-DRL1 (trunc)** | *GPD* promoter, N-tagged 3xV5 *DRL1* CD (2μ, *LEU*) | This Study |
| **pRS425-GPD-GFP-*ccdB*** | Destination vector: N-tagged GFP (2μ, *LEU*) | (Alberti et al., 2007) |
| **pRS425-GPD-GFP-DRL1** | *GPD* promoter, N-tagged GFP *DRL1* (2μ, *LEU*) | This study |
| **pRS425-GPD-GFP-DRL1 (CD)** | *GPD* promoter, N-tagged GFP *DRL1* CD (2μ, *LEU*) | This study |

**Table 3.**
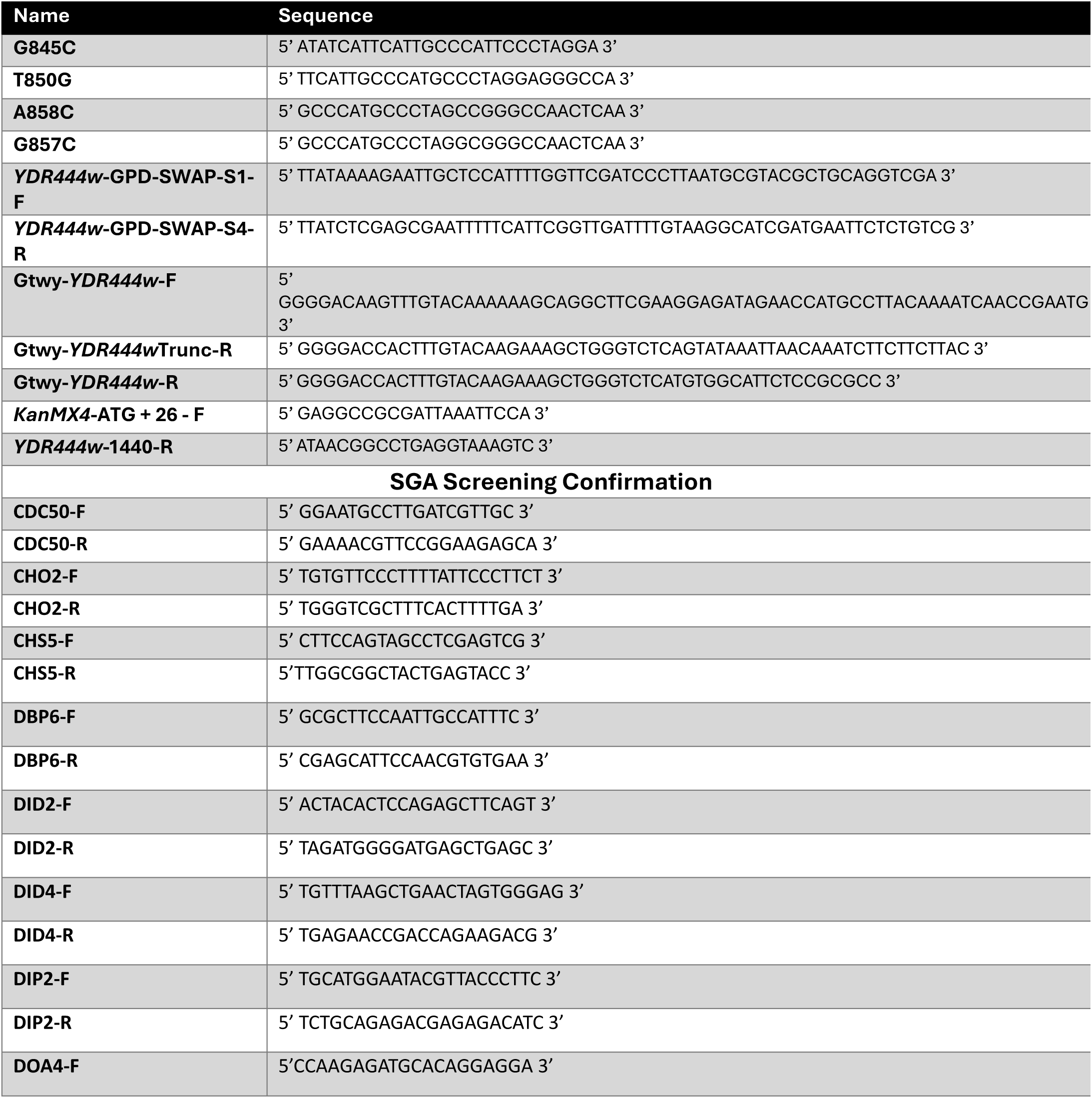
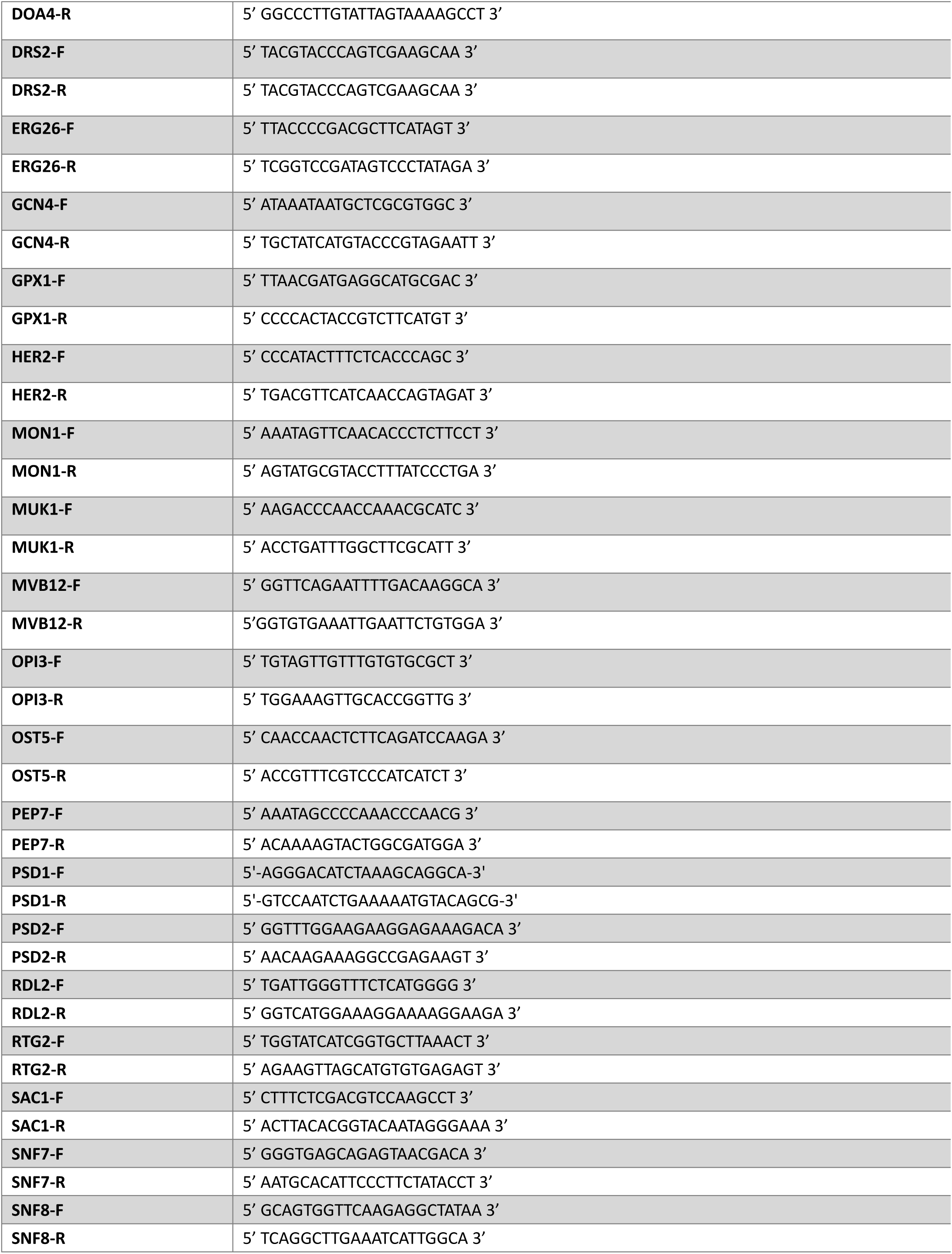
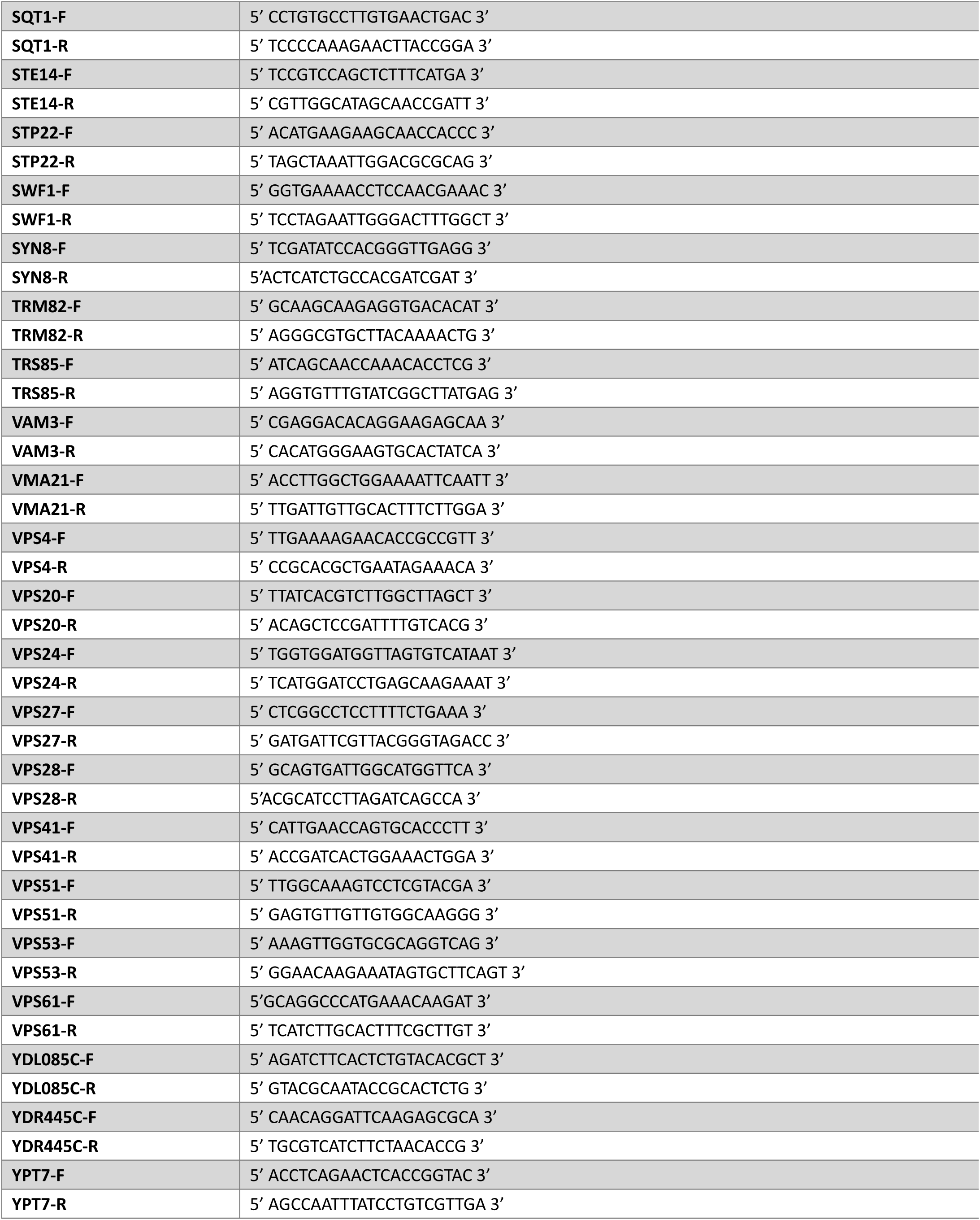
Primers used in this study.

### Growth

Yeast strains were grown in synthetic defined media (SD): 0.67% (w/v) yeast nitrogen base without amino acids and with ammonium sulphate, 2% (w/v) dextrose. Amino acids and nucleotides were supplemented based on growth requirements as follows: 0.002% (w/v) arginine, methionine, histidine, tryptophan, uracil and adenine sulfate, and 0.003% (w/v) leucine and lysine. Synthetic complete media (SC) refers to SD media supplemented with all the mentioned nutrients. Yeast extract peptone dextrose (YPD) media (2% peptone, 1% yeast extract, 2% dextrose) was used if only antibiotic selection was required. Solid media was made by adding 2% (w/v) agar.

Unless indicated, yeast cultures were grown at 30°C with shaking (200 rpm). Yeast transformations followed the Lithium Acetate method (Gietz et al., 1992). For genomic integrations, cells were grown for an additional 2 hours (30°C) to allow for homologous recombination to occur. All experiments were conducted in exponential growth phase. To achieve this, cells were grown overnight in the indicated growth media, then back diluted the next day to an OD_600_ of ∼0.2-0.3. The cells were incubated for 4-6 hours (30°C) after which OD_600_ was measured to ensure that the cells underwent a minimum of 1.5 cell divisions.

For cells carrying the pUG36-GFP-ALOD4 plasmid which contains a *MET25* promoter and is repressed by methionine, log phase cells were transferred to media lacking methionine and grown for an additional 30 minutes with shaking (30°C) to induce the GFP-ALOD4 probe expression prior to imaging.

### Yeast library construction for SGA analysis

A SGA query strain carrying the C1δ-GFP plasmid [*his3*Δ1 *leu2*Δ0 *met15*Δ0 *ura3*Δ0 *can1*Δ::*STE2*pr-*spHIS5 lyp1*Δ::*STE3*pr-*LEU2* Vam6-mCherry::NatR (pGPD416-C1δ-GFP)] was constructed and mated against the yeast deletion and DAmP libraries (Breslow et al., 2008; Giaever et al., 2002). The query strain has two mutations, *can1*Δ and *lyp1*Δ, that allow resistance against the toxic amino acid derivatives Canavanine and Thialysine. Arginine and lysine were omitted from the selection media to avoid competition for uptake with canavanine and lysine respectively during haploid selection. A RoToR bench-top colony arrayer (Singer Instruments) was used to manipulate libraries in high-density formats (1,536 array), where haploid strains from opposing mating types, each harboring a different genomic modification were mated on rich media plates. Diploids were selected on SD-Uracil plates containing Geneticin (Calbiochem-Merck) (200µg/mL) and Nourseothricin (Werner BioAgents) (200µg/mL). Sporulation was then initiated by shifting cells into nitrogen starvation media for 7 days. Haploid cells containing the desired deletion were selected by plating cells onto SD-Uracil plates containing Geneticin (200µg/mL) and Nourseothricin (200µg/mL), along with Canavanine and Thialysine (Sigma-Aldrich) for counterselection against remaining diploids, and lacking histidine to select for spores with an α mating type.

### Automated high-throughput fluorescence microscopy for the tailor-made library

The emerging yeast library containing the vacuolar markers, the DAG sensor and a deletion/hypomorphic allele in each yeast gene, was visualized using automated microscopy. Using the RoToR bench-top colony arrayer, cells were transferred from 1536-well plates into liquid media in 384-well plates and allowed to grow overnight in shaking incubator (LiCONiC Instruments) at 30°C in SD-Uracil. Cells were diluted to OD_600_∼0.2 using a JANUS liquid handler (Perkin Elmer) that is connected to the incubator and allowed to grow for 4 hours into mid-log phase. The cultures were then transferred into glass-bottom 384 well microscope plates (Matrical Bioscience) coated with Concanavalin A (Sigma-Aldrich) for 20 min to prevent cell movement during image acquisition. Wells were then washed twice to remove non-adherent cells and plates were then transferred to the ScanR automated inverted fluorescent microscope system (Olympus) using a Hamilton robotic swap arm. Following autofocus, images of cells were captured using a 60×air lens (NA 0.9) equipped with a cooled ORCA-ER charge-coupled device camera (Hamamatsu). Images were acquired at excitation filter 490/20 nm, emission filter 535/50 nm for GFP and at excitation filter 572/35 nm, emission filter 632/60 nm for mCherry. Three images were acquired for each strain in the library. Colony PCR was conducted to confirm the drastic hits from the SGA analysis using primers upstream and downstream of respective single gene deletion or insertions.

### Fluorescence Microscopy

Cells grown to log phase were concentrated and placed onto a 2% agarose pad comprised of the respective culture media. Images were acquired with a Zeiss Axio Imager Z2 upright epifluorescence microscope. ZEISS Zen blue imaging software and Zeiss plan Apochromat 100×/1.4 oil immersion objective lens was used for image acquisition. Colibri 7 LED light and 90 High Efficiency (HE) filter sets were used for excitation of GFP and mCherry. For GFP signal, samples were excited at 470/40 nm, and emission range was set at 525/50 nm while for mCherry signal, samples were excited at 555/30 nm, and emission range was set at 592/25 nm. For FM4-64 staining, 32 μM of the dye was added and incubated at 30°C for 45 minutes. The cells were then washed twice with their respective growth media and imaged immediately. For Edelfosine treatment, 20 μM of the drug or just vehicle (0.095% ethanol) was added followed by incubation at 30°C for 20 minutes. Cells were imaged immediately afterwards.

Z-stacks were taken in 10, 0.4 μm steps and deconvoluted in the ZEISS Zen blue imaging software using the constrained iterative method. Images were processed using the Fiji software for contrast adjustments, channel merging, batch cropping and scale bar addition. For clarity, the images shown are single planes.

### High resolution Airyscan Microscopy

Overnight cultures of WT (BY4741) cells simultaneously expressing GPD-C1δ-GFP and GPD-mCherry-ALOD4 were started in 3 mL SD-Leu-Ura. Cultures were diluted the next day to 0.2 OD_600_ in 2 mL SD-Leu-Ura and grown for 5 hours.. Cells were added to wells in a 96-well plate coated with Concanavalin A (Sigma-Aldrich) for 20 min to prevent cell movement during image acquisition. Wells were then washed twice to remove non-adherent cells and resuspended in SD-Leu-Ura medium containing edelfosine (20 μM) or vehicle control (0.095% ethanol) media. Cells were imaged at 10-minute intervals.

Image acquisition was performed using the Zeiss Celldiscoverer 7/7 Rev. 2 with Plan-Apochromat 50x/1.2 objective and 1x Tubelens optovar on Airyscan MPLX HS mode. Red channel was set to 5.00% laser power with 587nm/610nm excitation/emission wavelengths and 585-650 nm detection wavelength. Green channel was set to 3.00% laser power with 488nm/509nm excitation/emission wavelengths and 491-556 nm detection wavelength.

Images were 3D Airyscan processed and imported to Arivis Vision4D v4.1.1 (Zeiss) for analysis. Budding cells were selected for in-depth analysis, with 15 slices representing 2.66 μm (0.19 μm slice width) at the centre of the cell analysed. Sequentially, signal around the cell and interior signal from the cell was masked using the Object Mask function, leaving only signal attributable to probe-binding at the selected plasma membrane area. Intensity histograms for both channels were adjusted to ensure signal was not saturated at time zero. Objects based on signal from each channel at each time point were created using the Intensity Threshold Segmenter function (with the simple global thresholding method), ensuring that object boundaries and signal boundaries corresponded visually. Quantification of lipid domain volumes at each time point was done by comparing voxel count of objects created from either channel signal. The volume corresponding to either DAG or sterols probes was determined by adding the volume unique to that channel (i.e. excluding volume overlapping between both channels) and half of the volume overlapping between both channels. Total volume was determined by adding the volume from both channels such that volume overlapping between both channels was only accounted for once (i.e. volume unique to one channel + volume unique to other channel + volume overlapping between both channels).

### Subcellular Fractionation

Insoluble/ membrane fractions were prepared following a previously published protocol (Marr et al., 2012) with minor modifications. Mid-log phase *drl1*Δ cells (200 OD_600_) expressing pRS425-*GPDpr-GFP;* pRS425-*GPDpr-GFP-DRL1* or pRS425-*GPDpr-GFP-DRL1^CD^* were harvested by centrifugation, washed with distilled water, and resuspended in 0.7 mL of lysis buffer (20% glycerol, 50 mM Tris-HCl, 1 mM EDTA, pH 7.5, Complete EDTA-free protease inhibitor mixture [Roche], 1 mM PMSF, and 3 µg/mL pepstatin). Cells were lysed by adding 0.5 mm glass beads and vortexing on high speed (6 x 30 seconds) with a 30 second break on ice between vortexing steps. The beads were washed with 400 μL of lysis buffer, combined with the lysate, then spun at 1500 x g for 5 minutes at 4°C to remove cell debris. The cleared cell lysate was then ultracentrifuged at 450,000 x g for 15 minutes (4°C) using a Beckman Optima™ MAX ultracentrifuge equipped with a TLA 100.2 fixed angle rotor. The supernatant representing the soluble fraction was collected and the pellet was washed with 700 μL lysis buffer and ultracentrifuged again as previously described. The supernatant was then discarded and the pellet was resuspended in 0.5 mL lysis buffer with a dounce homogenizer (Cole-Parmer). Protein concentration of lysate, soluble and insoluble fractions were determined using the BCA assay (Thermo) and equal amount of protein per samples were analyzed by Western blot.

### Protein extraction and Western Blot

For analysis of cell lysates, 15-20 OD_600_ of cells were resuspended in 350 µL lysis buffer (20% glycerol, 50 mM Tris-HCl, 1 mM EDTA, pH 7.5, Complete EDTA-free protease inhibitor mixture [Roche], 1 mM PMSF, and 3 µg/mL pepstatin). Cells were lysed mechanically with 0.5 mm acid-washed glass beads using a Mini-Beadbeater (BioSpec) for 90 seconds. Lysates were centrifuged at 10,000 rpm (8,300 x g) for 1 minute at 4°C (Eppendorf 5415R), and supernatants were collected and saved.

Western Blot was carried out by the method of Laemmli (Laemmli, 1970). Briefly, proteins were separated by 10% resolving gel containing 2,2,2-trichloroethanol (TCE; Sigma) to visualize total protein loading (Ladner et al., 2004). Total protein content was imaged by excitation with trans UV light (302 nm) using a Bio-Rad Molecular Imager Gel Doc^TM^ XR. Proteins were transferred to a polyvinylidene fluoride (PVDF) membrane (Millipore) using a Bio-Rad Criterion Blotter at 30V for 16 hours and then stained with Red Ponceau (Sigma) to confirm transfer. Primary antibodies used were 1:2000 mouse αV5 (R96025; Thermo),1:2000 mouse αGFP (MA5-15256; Thermo). Horseradish peroxidase conjugated secondary antibodies (G21040; Thermo) were detected by chemiluminescence using ECL reagent (Amersham, GE Healthcare), and were imaged using an iBright FL1500 imager (Invitrogen).

Protein determination was carried out using a BCA protein assay kit (Pierce) with BSA as standard in GTE buffer (20% (v/v) glycerol, 50 mM Tris-HCl (pH 7.4), 1 mM EDTA (pH 8.0), adjusted to pH 7.5). Absorbance at 562 nm was recorded using a BioTek Synergy H1 microplate reader setup with SoftMax Pro software (Molecular Devices).

### Lipidomic Analysis

For lipidomic analysis, *KanMX-GPDpr-DRL1, BY4741* (*HO::KanMX*) and *drl1*Δ (*DRL1::KanMX*) cells were grown overnight in SC + 0.5 g/L G418 (Calbiochem). The cells were back diluted and grown overnight once more in SC. The next day, the cells were back diluted in SC and grown to mid log phase.

Extraction and analysis of lipids was performed as previously described in (Malitsky et al., 2016) with some modifications. Pellets containing 25 OD_600_ of cells were extracted with 1 ml of a pre-cooled (−20°C) homogenous methanol:methyl-tert-butyl-ether (MTBE) 1:3 (v/v) mixture, containing following internal standards: 0.1 μg/mL of Phosphatidylcholine (17:0/17:0) (Avanti), 0.4 μg/mL of Phosphatidylethanolamine (17:0/17:0, 0.15 nmol/mL of Ceramide/Sphingoid Internal Standard Mixture I (Avanti, LM6005), 00.4 μg/mL d5-TG Internal Standard Mixture I (Avanti, LM6000) and 0.1 μg/mL Palmitic acid-^13^C (Sigma, 605573). The tubes were vortexed and then sonicated for 30 min in ice-cold sonication bath (taken for a brief vortex every 10 minutes). Then, UPLC-grade water: methanol (3:1, v/v) solution (0.5 mL) was added to the tubes followed by centrifugation. The upper organic phase was transferred into a 2 mL Eppendorf tube. The polar phase was re-extracted as described above with 0.5 mL of MTBE. Both organic phases were combined and dried in speedvac and then stored at −80°C until analysis. For analysis, the dried lipid extracts were re-suspended in 200 μl mobile phase B (see below) and centrifuged again at 20800 x g (4°C) for 5 minutes.

Lipid extracts were analyzed using a Waters ACQUITY UPLC system coupled to a Vion IMS QTof mass spectrometer (Waters Corp., MA, USA). Chromatographic conditions were as described in (Malitsky et al., 2016) with small alterations. Briefly, the chromatographic separation was performed on an ACQUITY UPLC BEH C8 column (2.1×100 mm, i.d., 1.7 μm) (Waters Corp., MA, USA). The mobile phase A consisted of DDW: Acetonitrile: Isopropanol 46:38:16 (v/v/v) with 1% 1 M NH4Ac, 0.1% glacial acetic acid. Mobile phase B composition is DDW: Acetonitrile: Isopropanol 1:69:30 (v/v/v) with 1% 1 M NH4Ac, 0.1% glacial acetic acid. The column was maintained at 40°C; the flow rate of the mobile phase was 0.4 mL/minute, run time was 25 minutes. Linear gradient was as follows: Mobile phase A was run for 1 minute at 100%, then it was reduced to 25% for 11 minutes, followed by a decrease to 0% for 4 minutes. Then, mobile phase B was run at 100% for 5.5 min, following by setting mobile phase A to 100% for 0.5 minutes. Finally, the column was equilibrated at 100% A for 3 minutes. MS parameters were as follows: the source and de-solvation temperatures were maintained at 120°C and 450°C, respectively. The capillary voltage was set to 3.0 kV and 2 kV for positive and negative ionization mode, respectively; cone voltage was set for 40 V. Nitrogen was used as de-solvation gas and cone gas at a flow rate of 800 L/hr and 30 L/hr, respectively. The mass spectrometer was operated in full scan HDMSE resolution mode over a mass range of 50–2000 Da. For the high-energy scan function, a collision energy ramp of 20–80 eV was applied, for the low energy scan function – 4 eV was applied.

LC-MS data were analyzed and processed with UNIFI (Version 1.9.4, Waters Corp., MA, USA). The putative annotation of the lipid species was performed by comparison of accurate mass (below 5 ppm), fragmentation pattern, retention time (RT) and ion mobility (CCS) values to an in-house-generated lipid database. Peak intensities of the identified lipids were normalized to the internal standards and the amount of tissue used for analysis. For triplicates with one missing value, data was imputed as the mean of the two remaining replicates. For triplicates with two missing values, data was imputed based on the lowest normalized intensity for the given lipid species across all experimental conditions. Lipids that have not been described in the yeast lipidome were discarded. For lipid species with multiple isomers, we focused only on the most statistically significant isomer (lowest p-value). Lipids were considered to be significant if they satisfied: **1)** a fold change greater than 1.5 (log2(fold change) > 0.585); **2)** a p value less than 0.05 (-log10(pval) > 1.3). F-test was performed to ensure equal variance. If F-score < 0.05, a two-tailed unequal variance t-test was used, else a two-tailed equal t-test was used. Principal component analysis was performed in Metaboanalyst version 6.0 (Xia et al., 2009). Data was normalized using auto-scaling.

### AlphaFold 3 structure analysis

AlphaFold 3 structure prediction was performed using the publicly available AlphaFold Server (Abramson et al., 2024). Drl1 protein sequence was taken from Uniprot database (accession number Q04093) and built-in ligand selection was used. Default template settings were used for prediction. Molecular graphics and analyses performed with UCSF ChimeraX version 1.10.1, developed by the Resource for Biocomputing, Visualization, and Informatics at the University of California, San Francisco, with support from National Institutes of Health R01-GM129325 and the Office of Cyber Infrastructure and Computational Biology, National Institute of Allergy and Infectious Diseases (Meng et al., 2023).

### Molecular Docking

Structural models of Drl1 in complex with lysophospholipids were generated using Boltz-2, an open-source deep learning framework for biomolecular structure prediction (Passaro et al., 2025) (https://github.com/jwohlwend/boltz). LysoPC 18:1, the largest ligand considered, was used to generate the initial Drl1–ligand complex to ensure full accommodation of the binding pocket. Complexes with LysoPE and oleic acid (OLE) were subsequently derived by removing ligand-specific atoms from the LysoPC-bound structure, retaining only the atoms corresponding to each ligand of interest. All protein–ligand complexes were subjected to local energy minimization using the *Minimize Structure* tool in UCSF ChimeraX (https://www.cgl.ucsf.edu/chimerax/docs/user/tools/minimizestructure.html). Minimization was performed using the steepest descent algorithm with 100 steps. Binding free energies (ΔG_bind) and inhibition constants (Ki) were estimated using the AutoDock4 scoring function, as implemented within the Boltz-2 workflow. Protein–ligand contacts were analyzed by identifying residues within 5 Å of the ligand headgroup. Structural figures were generated using UCSF ChimeraX.

### Gene ontology analysis

Enrichment analysis was conducted using Cytoscape version 3.10.4 with ClueGO plugin version 2.5.10. The “Saccharomyces cerevisiae [559292, 4932]” marker list was loaded, and the list of 51 mutant genes was inputted with systematic names. GO Cellular Component or Biological Process ontologies were loaded and selected (enrichment analysis for each GO category was performed separately). The GO term/pathway selection option was adjusted to require a minimum of 3 genes, and the GO tree interval option was adjusted to allow for a minimum level of 0 and maximum level of 20.

### Statistical analysis

All statistical analysis was performed in GraphPad Prism 9.3.1. For numerical quantification analyzing the difference between only two variables, two-tailed nested t-tests were performed. For analysis of differences between WT and mutants nested one-way ANOVA with Šídák’s multiple comparisons pos-test were performed.

### Supplementary Data

**Table S1**. List of deletion and DAmP library hits

**Table S2**. Lipidomics data

**Figure S1**. Mutants Exhibiting Cell Periphery Phenotype.

**Figure S2**. Drl1 and Rog1 3D catalytic domain alignment.

**Figure S3**. Drl1 does not alter cytosolic facing pools of PA, PS, PI(4)P or PI(4,5)P_2_ -

**Figure S4**. Drl1 modulates fatty acid and lysolipid levels.

---

**Figure S1.**
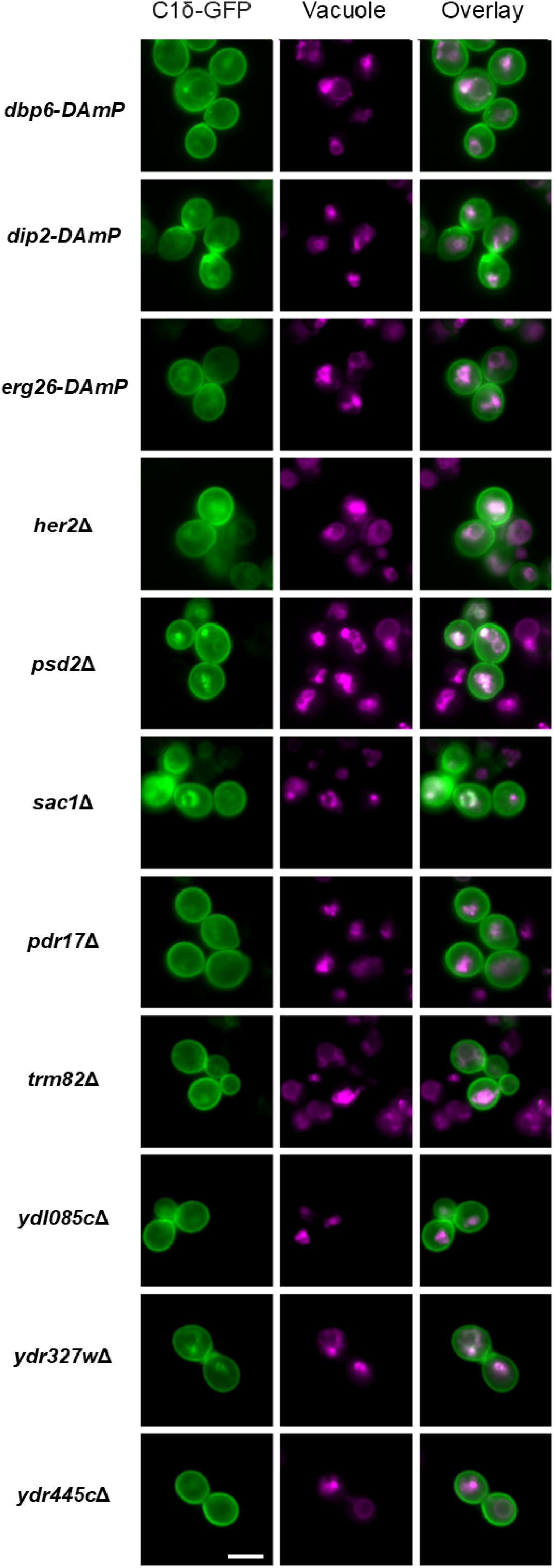
Mutants Exhibiting Cell Periphery Phenotype. Images captured from the high throughput genetic screen for the “cell periphery” category including 5 non-essential genes (*PSD2, SFH4, SAC1, TRM82, HER2),* 2 essential genes (*DBP6, ERG26) and 3 unknown non-essential genes (YDR445C, YDR327W, YDL085C).* Diacylglycerol is visualized by the C1δ-GFP lipid probe, and the vacuole is visualized with Vam6-mCherry. Scale bar 5 µm.

**Figure S2:**
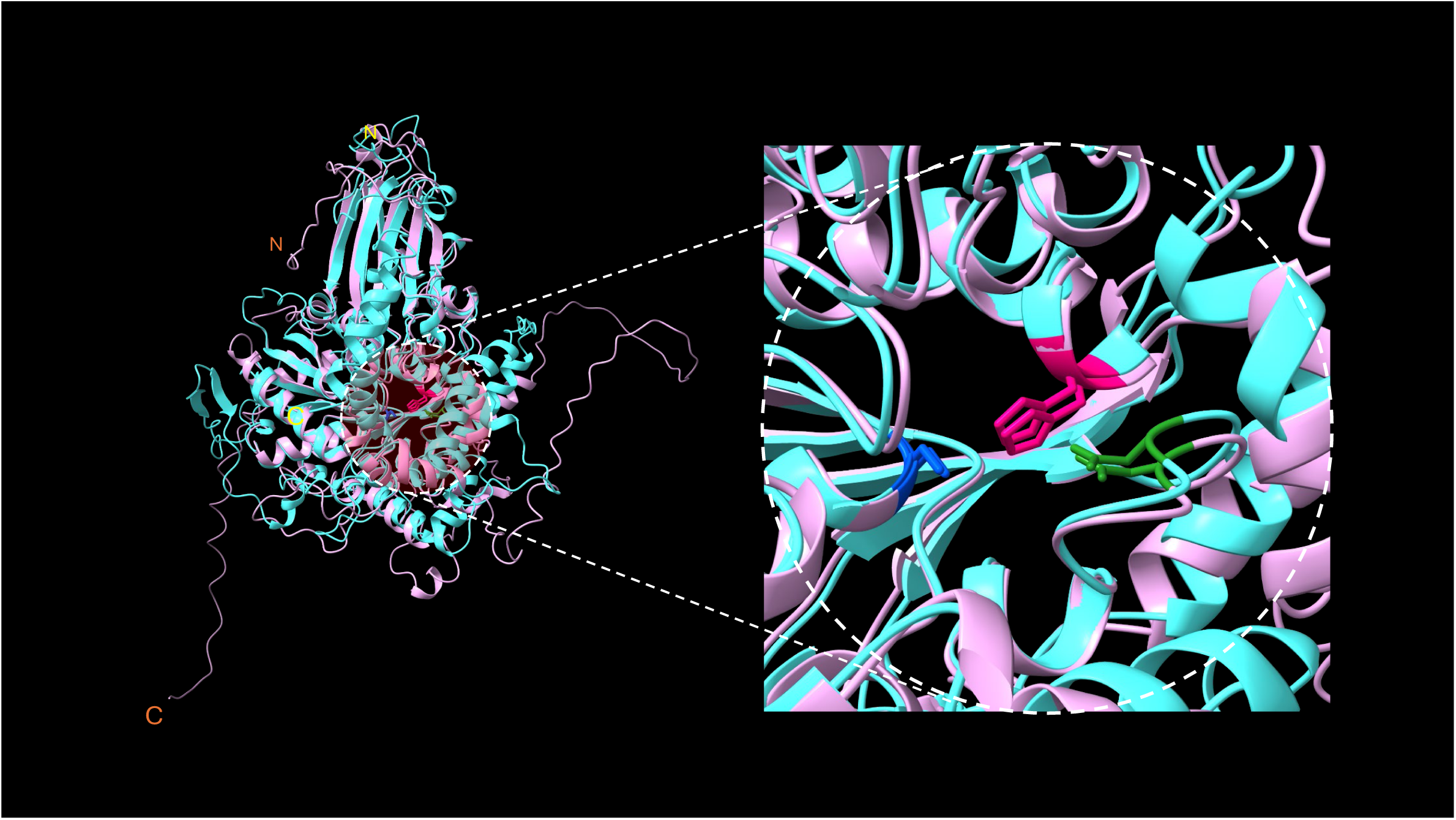
Drl1 and Rog1 3D catalytic domain alignment. Aligned predicted structures for Drl1 and Rog1 highlighting conservation of Ser-His-Asp catalytic triad in 3-dimensional space. Drl1 is coloured in light pink, Rog1 in light blue. Catalytic serine (284 in Drl1, 269 in Rog1) is coloured in pink, histidine (617 in Drl1, 552 in Rog1) is coloured in dark blue, aspartate (405 in Drl1, 372 in Rog1) is coloured in green. Drl1 N- and C-terminus are labeled in orange, Rog1 N- and C-terminus are labeled in yellow.

**Figure S3.**
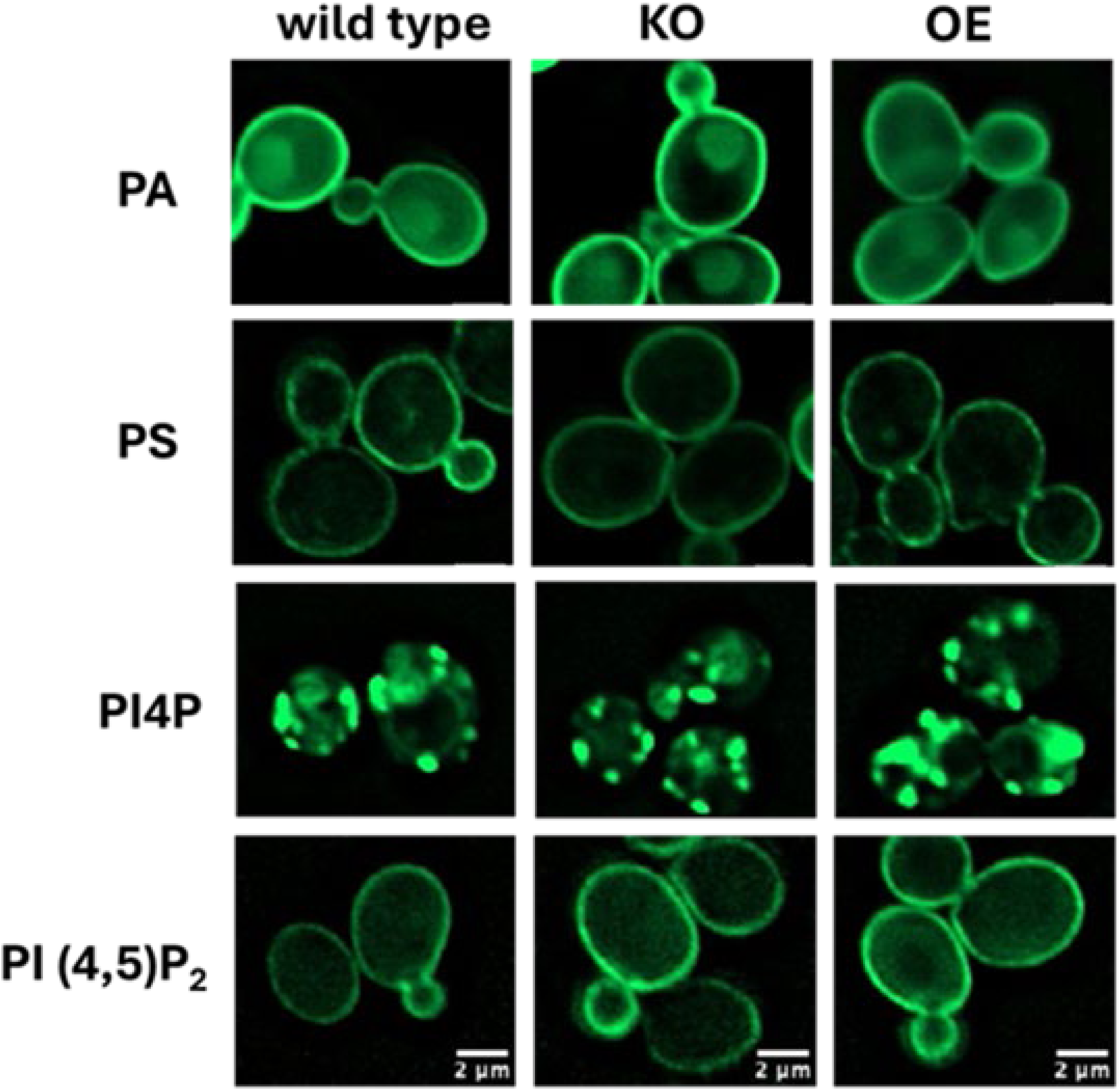
Drl1 does not alter cytosolic facing pools of PA, PS, PI(4)P or PI(4,5)P_2_. Wild-type (WT), *drl1*Δ (KO) and *DRL1* overexpressing (OE) cells were transformed with plasmids to express probes for detection of cytosolic facing pools of PA (GFP-Spo20), PS (GFP-Lact-C2), PI(4)P (GFP-PH-FAPP1) and PI(4,5)P_2_ (GFP-2xPH-PLCδ). Transformants were grown on selective defined medium to log phase followed by live imaging as indicated in M&M. Scale bar = 2 μm.

**Figure S4.**
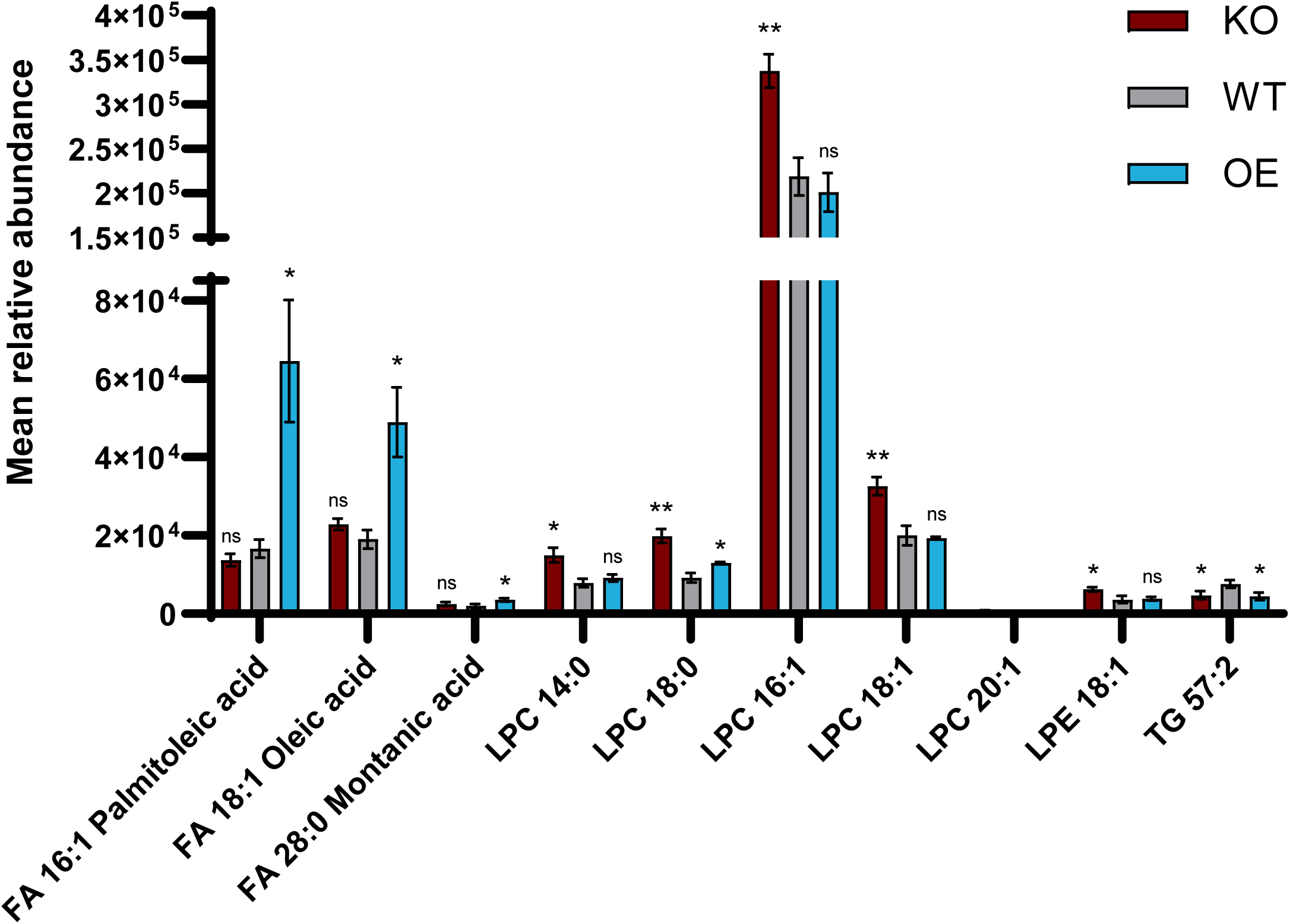
Drl1 modulates fatty acid and lysolipid levels. Mean relative abundance of significantly altered lipids among *drl1*Δ cells (KO), *DRL1* overexpression cells (OE) or WT BY4741 cells. (WT). +/- bars indicate standard deviation. ns = p>0.05, (*) = p<0.05, (**) = p<0.01.

