## Supplementary figures and images for "A Genome-wide Visual Screen Identifies Lysophosphatidylcholine as Counter Spatial Regulator of DAG and Sterols in Yeast"

### Figure S1

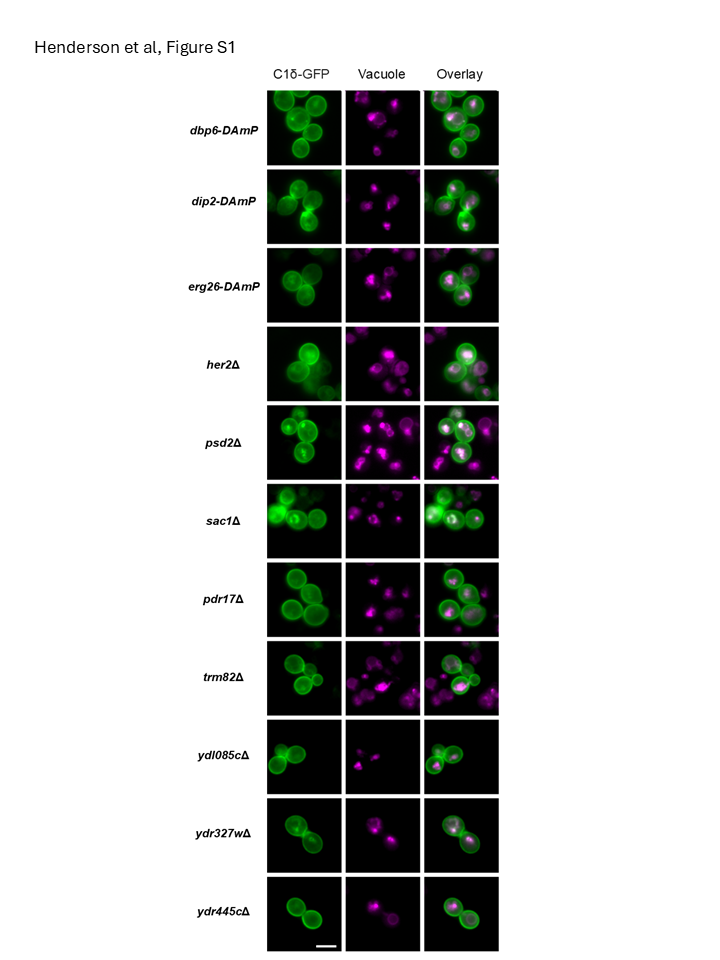

### Figure S2

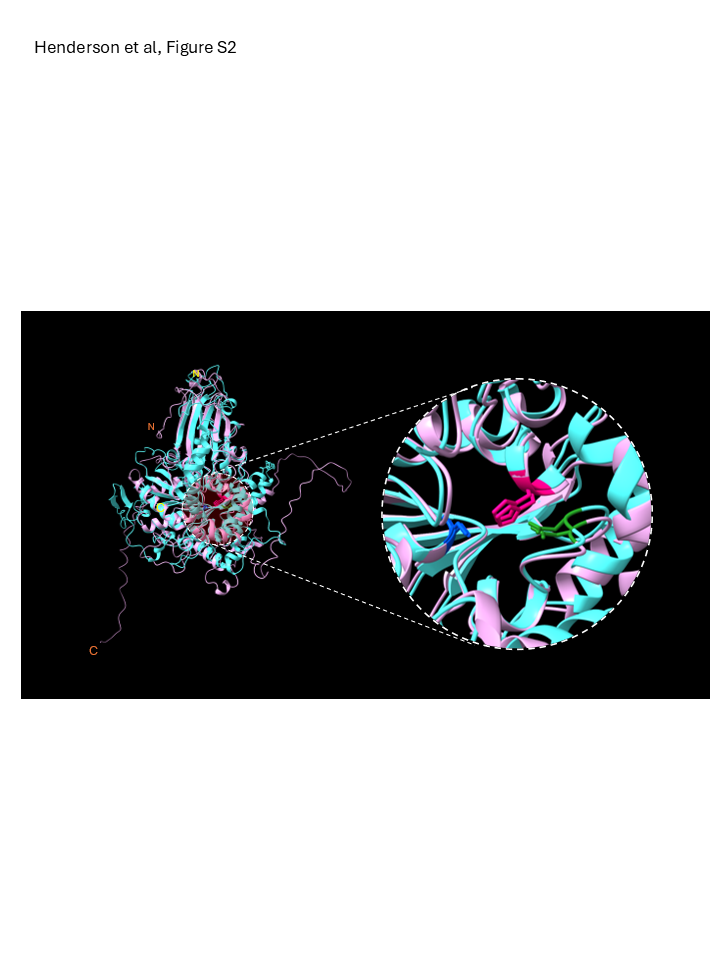

### Figure S3

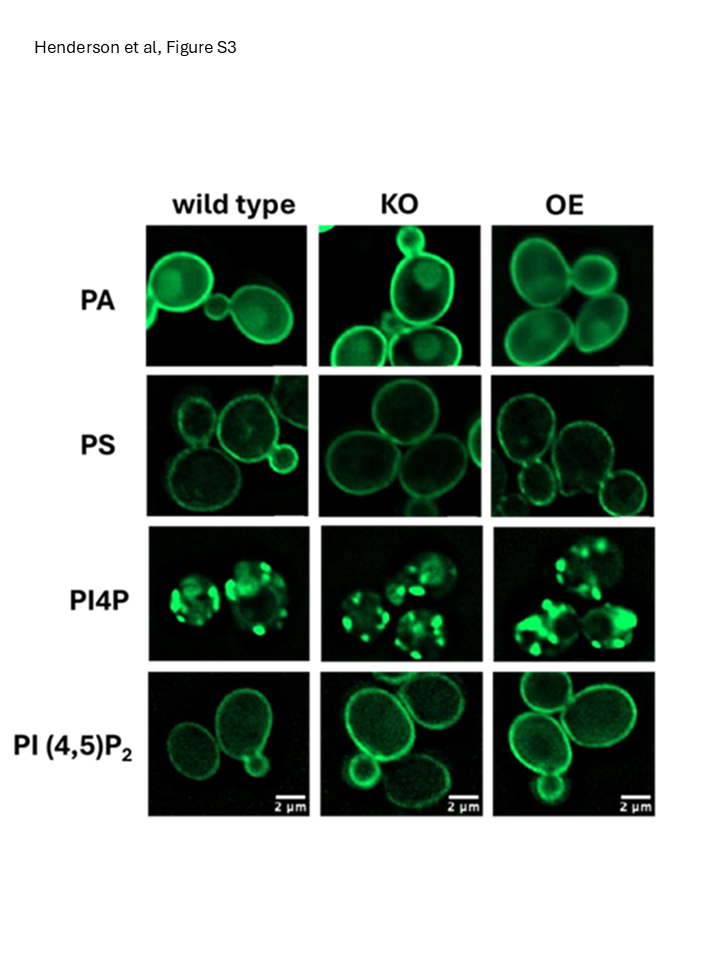

### Figure S4

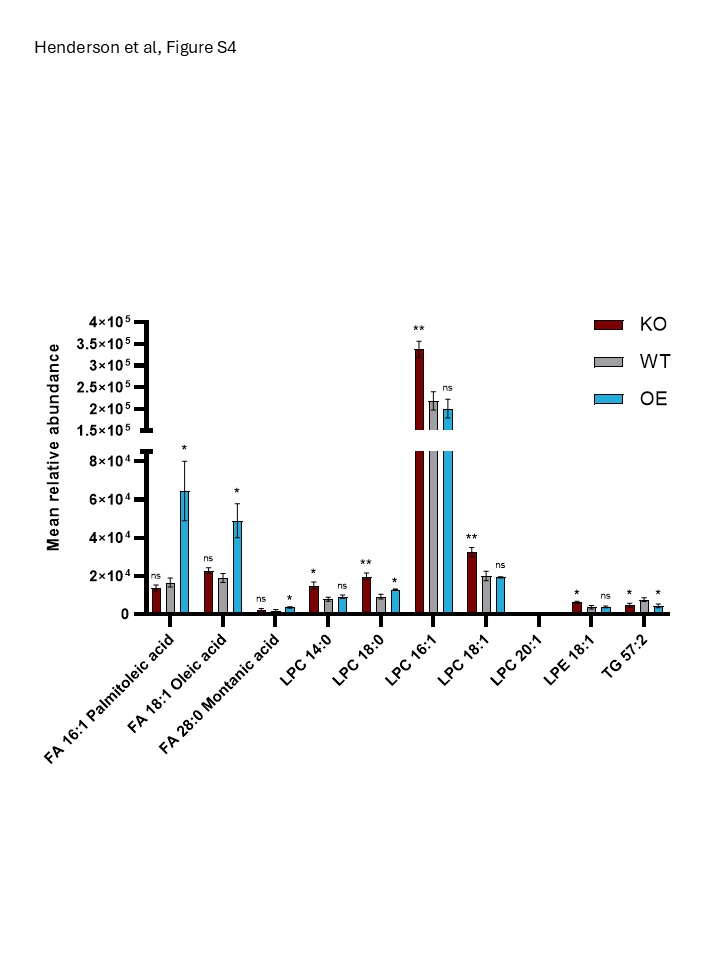
