## Supplementary material for "A Genome-wide Visual Screen Identifies Lysophosphatidylcholine as Counter Spatial Regulator of DAG and Sterols in Yeast": Table S1

**Supplementary Table S1.** List of deletion and DAmP library hits, their biological function and DAG sensor localization. PCR validation of the genotype is indicated with a "+"

| Systematic name | Standard name | Library | Cellular function (SGD) | PCR Val | C1δ-GFP localization |
|---|---|---|---|---|---|
| *YNR038W* | *DBP6* | DAmP | Essential protein involved in ribosome biogenesis | + | Cell Periphery |
| *YLR129W* | *DIP2* | DAmP | Nucleolar protein, required for 18S rRNA biogenesis | + | Cell Periphery |
| *YGL001C* | *ERG26* | DAmP | C-3 sterol dehydrogenase involved in ergosterol biosynthesis | + | Cell Periphery |
| *YMR293C* | *HER2* | Deletion | Subunit of the trimeric GatFAB AmidoTransferase(AdT) complex | + | Cell Periphery |
| *YGR170W* | *PSD2* | Deletion | Phosphatidylserine decarboxylase of the Golgi and vacuolar membranes | + | Cell Periphery |
| *YKL212W* | *SAC1* | Deletion | Phosphatidylinositol phosphate phosphatase | + | Cell Periphery |
| *YNL264C* | *PDR17* | Deletion | Phosphatidylinositol transfer protein (PITP); downregulates Plb1p-mediated turnover of phosphatidylcholine; forms a complex with Psd2p which appears essential for maintenance of vacuolar PE levels | + | Cell Periphery |
| *YDR165W* | *TRM82* | Deletion | Noncatalytic subunit of a tRNA methyltransferase complex | + | Cell Periphery |
| *YDL085c* | *YDL085c* | Deletion | Homolog of modifier of aggregation-4 (MOAG-4) from worm and small EDRK-rich factor protein (SERF1a) from human; highly conserved enhancer of amyloid formation of Aβ40 and α-synuclein both in vitro and in vivo | + | Cell Periphery |
| *YDR327w* | *YDR327w* | DAmP | Unknown- Overlaps with *SKP1* | - | Cell Periphery |
| *YDR445c* | *YDR445c* | Deletion | Unknown- Named *DRL1* (this study) | + | Cell Periphery |
| *YGL124C* | *MON1* | Deletion | Membrane tethering and fusion | - | Clump/unique |
| *YPL070W* | *MUK1* | Deletion | Guanine nucleotide exchange factor (GEF) | - | Clump/unique |
| *YJR073C* | *OPI3* | Deletion | Methylene-fatty-acyl-phospholipid synthase | + | Clump/unique |
| *YNL169C* | *PSD1* | Deletion | Phosphatidylserine decarboxylase of the mitochondrial inner membrane | + | Clump/unique |
| *YDR126W* | *SWF1* | Deletion | Palmitoyltransferase that acts on transmembrane proteins | + | Clump/unique |
| *YAL014C* | *SYN8* | Deletion | Endosomal SNARE related to mammalian syntaxin 8 | + | Clump/unique |
| *YOR106W* | *VAM3* | Deletion | Syntaxin-like vacuolar t-SNARE | + | Clump/unique |
| *YGR105W* | *VMA21* | Deletion | Vacuolar ATPase assembly | + | Clump/unique |
| *YDR080W* | *VPS41* | Deletion | Subunit of the HOPS endocytic tethering complex, membrane fusion | + | Clump/unique |
| *YML001W* | *YPT7* | Deletion | Rab family GTPase required for homotypic fusion event in vacuole inheritance | + | Clump/unique |
| *YCR094W* | *CDC50* | Deletion | Endosomal protein that interacts with phospholipid flippase Drs2p | + | Puncta |
| *YGR157W* | *CHO2* | Deletion | Phosphatidylethanolamine methyltransferase (PEMT) | + | Puncta |
| *YKR035W-A* | *DID2* | Deletion | Class E protein of the vacuolar protein-sorting (Vps) pathway | - | Puncta |
| *YKL002W* | *DID4* | Deletion | Class E Vps protein of the ESCRT-III complex | - | Puncta |
| *YDR069C* | *DOA4* | Deletion | Endocytosis and ubiquitin homeostasis | + | Puncta |
| *YAL026C* | *DRS2* | Deletion | Trans-golgi network aminophospholipid translocase (flippase) | + | Puncta |
| *YEL009C* | *GCN4* | Deletion | transcriptional activator of amino acid biosynthetic genes | + | Puncta |
| *YGR206W* | *MVB12* | Deletion | ESCRT-I subunit required to stabilize ESCRT-I core complex oligomers | + | Puncta |
| *YGL226C-A* | *OST5* | Deletion | N-linked glycosylation | + | Puncta |
| *YDR323C* | *PEP7* | Deletion | Adaptor protein involved in vesicle-mediated vacuolar protein sorting | + | Puncta |
| *YGL252C* | *RTG2* | Deletion | Transcription | - | Puncta |
| *YLR025W* | *SNF7* | Deletion | One of four subunits of the ESCRT-III complex | - | Puncta |
| *YPL002C* | *SNF8* | Deletion | Component of the ESCRT-II complex | + | Puncta |
| *YIR012W* | *SQT1* | DAmP | Specific chaperone for ribosomal protein Rpl10p | + | Puncta |

| | | | | | |
|---|---|---|---|---|---|
| *YCL008C* | *STP22* | Deletion | Component of the ESCRT-I complex | + | Puncta |
| *YDR108W* | *TRS85* | Deletion | Component of transport protein particle (TRAPP) complex III | + | Puncta |
| *YPR173C* | *VPS4* | Deletion | AAA-ATPase involved in multivesicular body (MVB) protein sorting | + | Puncta |
| *YMR077C* | *VPS20* | Deletion | Myristoylated subunit of the ESCRT-III complex | + | Puncta |
| *YKL041W* | *VPS24* | Deletion | One of four subunits of the ESCRT-III complex | + | Puncta |
| *YNR006W* | *VPS27* | Deletion | Ubiquitin binding protein involved in endosomal protein sorting | + | Puncta |
| *YPL065W* | *VPS28* | Deletion | Component of the ESCRT-I complex | + | Puncta |
| *YKR020W* | *VPS51* | Deletion | Component of the GARP (Golgi-associated retrograde protein) complex | + | Puncta |
| *YJL029C* | *VPS53* | Deletion | Component of the GARP (Golgi-associated retrograde protein) complex | + | Puncta |
| *YDR136C* | *VPS61* | Deletion | Protein targeting to vacuole | + | Puncta |
| *YPL078C* | *ATP4* | Deletion | Subunit b of the stator stalk of mitochondrial F1F0 ATP synthase | - | Soluble |
| *YLR330W* | *CHS5* | Deletion | Component of the exomer complex | + | Soluble |
| *YKL026C* | *GPX1* | Deletion | Phospholipid hydroperoxide glutathione peroxidase | + | Soluble |
| *YOR286W* | *RDL2* | Deletion | Protein with rhodanese activity | + | Soluble |
| *YDR410C* | *STE14* | Deletion | Protein methylation | - | Soluble |
| *YGR020C* | *VMA7* | Deletion | Vacuolar ATPase | - | Soluble |
